# Dynamics-aware Evolutionary Profiling Uncouples Structural Rigidity from Functional Motion to Enable Enhanced Variant Interpretation

**DOI:** 10.64898/2026.01.16.699945

**Authors:** Taner Karagöl, Alper Karagöl

## Abstract

Evolutionary conservation is a powerful part of mutational intolerance prediction, yet traditional pathogenicity metrics frequently conflate two distinct biophysical constraints: structural stability (rigidity) and functional mechanics (dynamics). We introduce Dynamics-Aware Evolutionary Profiling to resolve this ambiguity, integrating Molecular Dynamics with evolutionary conservation and coupling analysis across human/cross-species-proteome of 151 protein structures. By mathematically uncoupling biophysical forces, we define orthogonal metrics; the <u>R</u>igid <u>C</u>onserved <u>S</u>core (RCS) for the structural scaffold, and the <u>D</u>ynamic <u>C</u>onserved <u>S</u>core (DCS) for flexible residues. Our analysis reveals a fundamental bifurcation in pathogenicity. RCS serves as a filter for lethal structural failure, isolating hydrophobic core residues whose mutation triggers unfolding. In contrast, DCS identified a rare population of residues that are evolutionarily highly-conserved but structurally mobile; these Dynamic-Conserved sites exhibit intermediate pathogenicity and are enriched in flexible hinge residues (Gly, Pro). Validation against 737 human variants from ClinVar demonstrates that DCS captures a distinct pathogenic mechanism regarding essential protein motion. Notably, DCS and RCS correctly flagged some pathogenic variants of NARS1 and PGK1 that were misclassified as benign or ambiguous by AlphaMissense. These results indicate that while the rigid core represents a stability bottleneck, DCS isolates functional sites likely driving allosteric regulation. We provide an open-access web interface (*ADEPT*) for these metrics. By isolating dynamic-conserved residues, this framework refines the interpretation of Variants of Uncertain Significance in dynamic regions and reveals tunable targets for rational drug design, moving beyond the static optimization of the folded state.

## Introduction

Traditionally, evolutionary conservation and pathogenicity have been interpreted mainly in terms of fold stability, with the prevailing view that strongly conserved residues form a rigid framework, and misfolding variants are causing pathogenicity [1, 2, 3]. However, while foundational, this perspective frequently oversimplifies the complex reality of protein biology by treating macromolecules as static entities. In reality, proteins are dynamic molecular machines [4, 5] whose biological function is defined not merely by their ability to fold, but by their capacity to execute specific atomic motions. Essential biological processes rely on specific residues that must remain intrinsically flexible to facilitate conformational transitions [6]. Paradoxically, these residues rarely may be also conserved [4, 6, 7] as if they constituted a rigid core, not for maintaining a static structure, but for ensuring accurate functional motions.

This requirement creates a complex-evolutionary landscape where proteins must satisfy competing biophysical constraints; the need for structural rigidity [1] to prevent unfolding, and the need for kinetic flexibility to enable function [6]. As result, evolutionary conservation reflects two chemically and mechanically distinct signals. One corresponds to the thermodynamic core, where mutations are selected against due to their destabilizing effects on the folded state [2]. The other, more subtle signal originates from the residues that function as allosteric switches or molecular hinges. Furthermore, certain residue substitutions may accommodate fold-preserving substitutions with minimal pathogenic impact, as observed for chemically distinct *QTY-code* substitutions [8, 9, 10, 11]. On the other hand, the substitutions in flexible regions at critical dynamic sites [4] or binding sites [12] may induce catastrophic functional pathogenicity without triggering global protein misfolding, reflecting a mechanism distinct from classical destabilization. Distinguishing scaffold-maintaining residues from those dynamic residues that drive functional flexibility remains a fundamental challenge in structural biology.

Despite the critical importance of these dynamic sites, current computational pathogenicity and evolutionary prediction algorithms struggle to identify them, and the structural information on proteins limited [3]. Recent improvements on protein structural prediction tools including AlphaFold3 [13] has enabled a revolution in modelling the structure of proteins. While state-of-the-art deep learning models, including AlphaMissense [14], have achieved remarkable success in predicting pathogenicity by learning the rules of structural stability from static crystal structures and multiple sequence alignments. However, these models often falter when resolving the functional nuances of flexible regions, sometimes misclassifying essential sites as ambiguous or benign. This limitation can arise because static predictors inherently bias their scoring towards packing density and rigidity, or do not use protein dynamics as a factor. As a result, a significant class of pathogenic variants, that can be classified as those that disrupt the dimension of time rather than the dimension of space, remains in the blind spots of proteomic analysis.

Recent computational advances have begun to combine evolutionary analysis with <u>m</u>olecular <u>d</u>ynamics (MD), notably through methods that map coevolutionary couplings onto dynamic trajectories to identify allosteric networks and functional residue pairs [15, 16, 17]. While these approaches successfully show correlated motions within protein communities, they typically treat dynamics and evolution as overlapping filters to identify sites of communication. Our approach differs by treating these forces as independent components rather than overlapping filters. By deconvoluting the evolutionary signal at the single-residue level, we distinguish conservation required for structural folding from conservation required for structural dynamics, thereby revealing a previously obscured dimension of molecular evolution. Unlike network-based methods that identify broad functional connectivity, our framework allows us to pinpoint cryptic variants in flexible regions that lack the structural rigidity to be flagged by standard predictors, information that is equally vital for rational protein engineering and molecular evolution.

To bridge the gap between rigid structure and dynamic function, we introduce <u>D</u>ynamics-aware <u>E</u>volutionary <u>P</u>rofiling (DEP), a quantitative framework that integrates molecular dynamics trajectories with deep evolutionary analysis. By mathematically uncoupling the biophysical forces of rigidity from dynamics, we establish a suite of orthogonal metrics designed to capture distinct biophysical states. We define the <u>R</u>igid <u>C</u>onserved <u>S</u>core (RCS) to identify the residues that are strictly conserved and structurally immobile. In parallel, we introduce the <u>D</u>ynamic <u>C</u>onserved <u>S</u>core (DCS) to isolate evolutionarily constrained flexible sites. This approach allows for a high-resolution dissection of the protein evolutionary landscape, moving beyond simple classifications of conserved versus variable to a multi-dimensional view that accounts for the specific dynamical role of each residue. We validated this framework using a *pan-proteome* dataset totaling over 30,000 residues, combining 93 *cross-species* alpha-helical proteins and 58 medium-length *homo sapiens* proteins. Our analysis demonstrates that RCS and DCS capture statistically independent information channels, successfully disentangling the conserved dynamics of the protein from its rigid scaffold. To allow broader community adoption and ensure reproducibility, we have developed an open-access web server, *ADEPT* (<u>A</u>utomated <u>D</u>ynamics-aware <u>E</u>volutionary <u>P</u>rofiling <u>T</u>ool) (https://www.karagolresearch.com/adept) that automates the calculation of these scores from standard input data.

Furthermore, this framework extends beyond individual residues to interrogate the role of “dynamic-conservation” as a mechanism for mutational tolerance. Unlike independent flexible loops which may be functionally neutral, coupled dynamic networks often facilitate long-range communication across protein domains. Understanding these dynamic-evolutionary relationships provides critical insights into the mechanisms of complex disease and evolutionary history, distinguishing between variants of stability (high RCS) and those driven by specific functional dynamics (high DCS). Additionally, we demonstrate the clinical utility of this framework by mapping genetic variants from the ClinVar database [18], revealing that DCS and RCS can also isolate cryptic pathogenic events that are misclassified as benign by predictors like AlphaMissense. By uncovering the interplay between structural rigidity, dynamic flexibility, and evolutionary conservation; this study offers a comprehensive map of residue evolution, challenging the dogma that conservation implies rigidity and offering a refined strategy for interpreting <u>V</u>ariants of <u>U</u>ncertain <u>S</u>ignificance (VUS) in rational drug design and precision medicine.

## Results

### Descriptive Statistics of the Biophysical Landscape

Our analysis encompassed a comprehensive pan-proteome dataset, derived from a *cross-species* alignment of 93 alpha-helical proteins and a targeted cohort of 58 medium-length *homo sapiens* (human) proteins (400-600 amino acids) (methods) from selected from available molecular dynamics trajectories ATLAS database [19] (Supplementary Table 1). To quantify the interplay between stability and motion, we defined two primary metrics; the <u>R</u>igid <u>C</u>onserved <u>S</u>core (RCS), calculated as the product of conservation and structural rigidity:

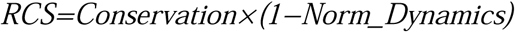

and the Dynamic Conserved Score (DCS), calculated as the product of conservation and normalized flexibility:

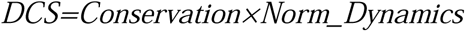

Furthermore, to capture the effects driving evolutionary communication, we defined the <u>D</u>ynamics and <u>R</u>igid <u>Co</u>-evolutionary Cou<u>p</u>ling <u>S</u>core (*DCopS* and *RCopS*) by integrating pairwise evolutionary couplings with dynamic fluctuation (analyzed in detail in subsequent sections). The evolutionary conservation scores and pairwise evolutionary couplings calculated from EVcouplings server [20]. The biophysical landscape of this combined dataset is characterized by a distributional asymmetry between evolutionary conservation and structural dynamics. The RCS, which targets the structural rigidity, exhibits a high baseline across the proteome (Human Mean μ≈4.43; Median ≈3.97), reflecting the pervasive requirement for thermodynamic stability in protein folding. In contrast, the DCS isolates a significantly sparser, heavy-tailed population (Human Mean μ≈1.14; Median ≈0.62). The mean scores of proteins from available molecular dynamics data is also calculated (Supplementary Figures 1, 2, 3, and 4).

While the vast majority of residues contribute to the dynamic background, high-scoring DCS residues represent very rare portions (Figure 1). Specifically, the 99th percentile of DCS values reaches 8.28, compared to a 95th percentile of 4.11, marking the top 1% as highly dynamic-conserved residues that rise sharply above the noise of the proteome. This distribution confirms that while structural rigidity is a global constraint widely distributed across the sequence, conserved flexibility is a localized, high-value feature.

**Figure 1.**
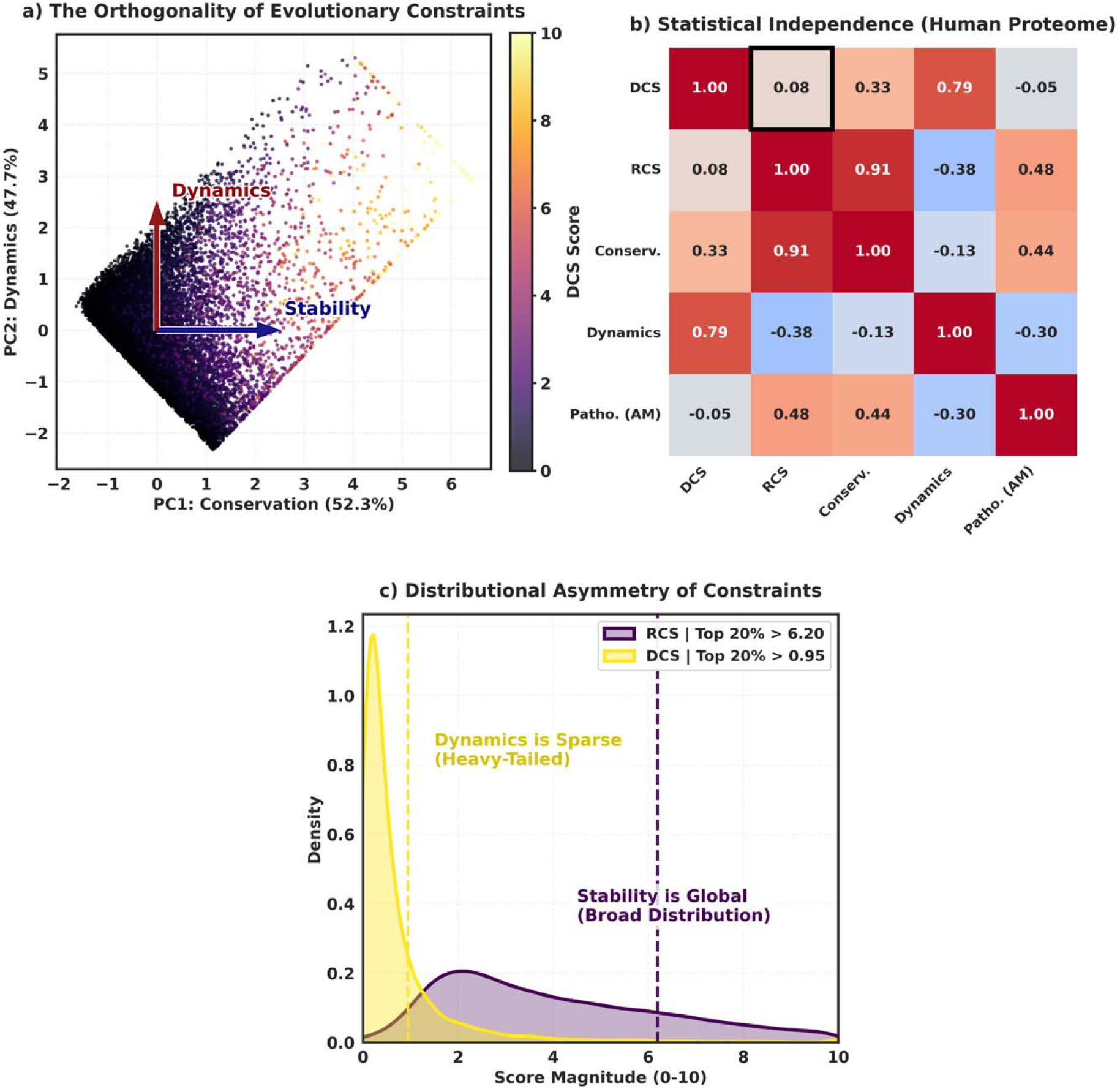
The Orthogonality of Stability and Motion. **(a)** <u>P</u>rincipal <u>C</u>omponent <u>A</u>nalysis (PCA) biplot of the pan-proteome residue landscape (93 alpha-helical proteins), illustrating the relationship between Evolutionary Conservation and Structural Dynamics. The two principal components (PC1, PC2) capture distinct axes of variation. **(b)** Spearman correlation matrix quantifying the relationships between derived metrics. <u>R</u>igid <u>C</u>onserved <u>S</u>core (RCS) exhibits a strong positive correlation with conservation (ρ=0.91), while the <u>D</u>ynamic <u>C</u>onserved <u>S</u>core (DCS) is statistically independent (ρ=0.08). **(c)** Density plots of DCS (orange) and RCS (purple) scores derived from the pan-proteome dataset, revealing a fundamental distributional asymmetry.

### Orthogonality of Stability and Motion

Unsupervised <u>P</u>rincipal <u>C</u>omponent <u>A</u>nalysis (PCA) [21] of the global dataset reveals a fundamental bifurcation in the evolutionary landscape. The variance in biophysical properties is partitioned nearly evenly between the first two principal components (PC1:52.3%, PC2: 47.7%) (Figure 1). This equipartition demonstrates that Evolutionary Conservation (loading primarily on PC1) and Structural Dynamics (loading on PC2) operate as mathematically orthogonal forces rather than redundant features.

This independence is quantitatively validated by Spearman correlation analysis, which shows a negligible correlation between the Dynamic Conserved Score (DCS) and the Rigid Conserved Score (RCS) (ρ≈0.08). Conversely, RCS exhibits a near-perfect correlation with raw conservation (ρ≈0.91) (Figure 1). This statistical dissociation validates our framework’s ability to uncouple the two primary drivers of protein constraint, stability and motion. It proves that traditional static conservation analysis captures only one half of the biophysical constraints shaping the proteome (the structural scaffold), while the orthogonal axis of functional dynamics remains largely invisible to standard evolutionary metrics.

### Mechanical Code for Dynamic Conservation

We have analyzed the propensity of specific amino acids to populate high-scoring regions. For the DCS, the Kruskal-Wallis test [22] confirmed that high DCS values are driven by a specific subset of residues rather than a random sampling of the sequence (Figure 2). Specifically, the heatmap highlights Glycine (Gly) (Propensity Z=1.77) and Proline (Pro) (Z=0.89 to 1.25). This can also provide a clear biophysical rationale; Glycine, lacking a side chain, allows for a wider range of dihedral angles essential for hinge bending, while Proline’s cyclic structure introduces intrinsic kinks that function as molecular pivots or switches [23]. Additionally, Methionine (Z=2.14), often acting as a flexible hydrophobic connections [24] may act as a key stabilizer of these dynamic interfaces.

**Figure 2.**
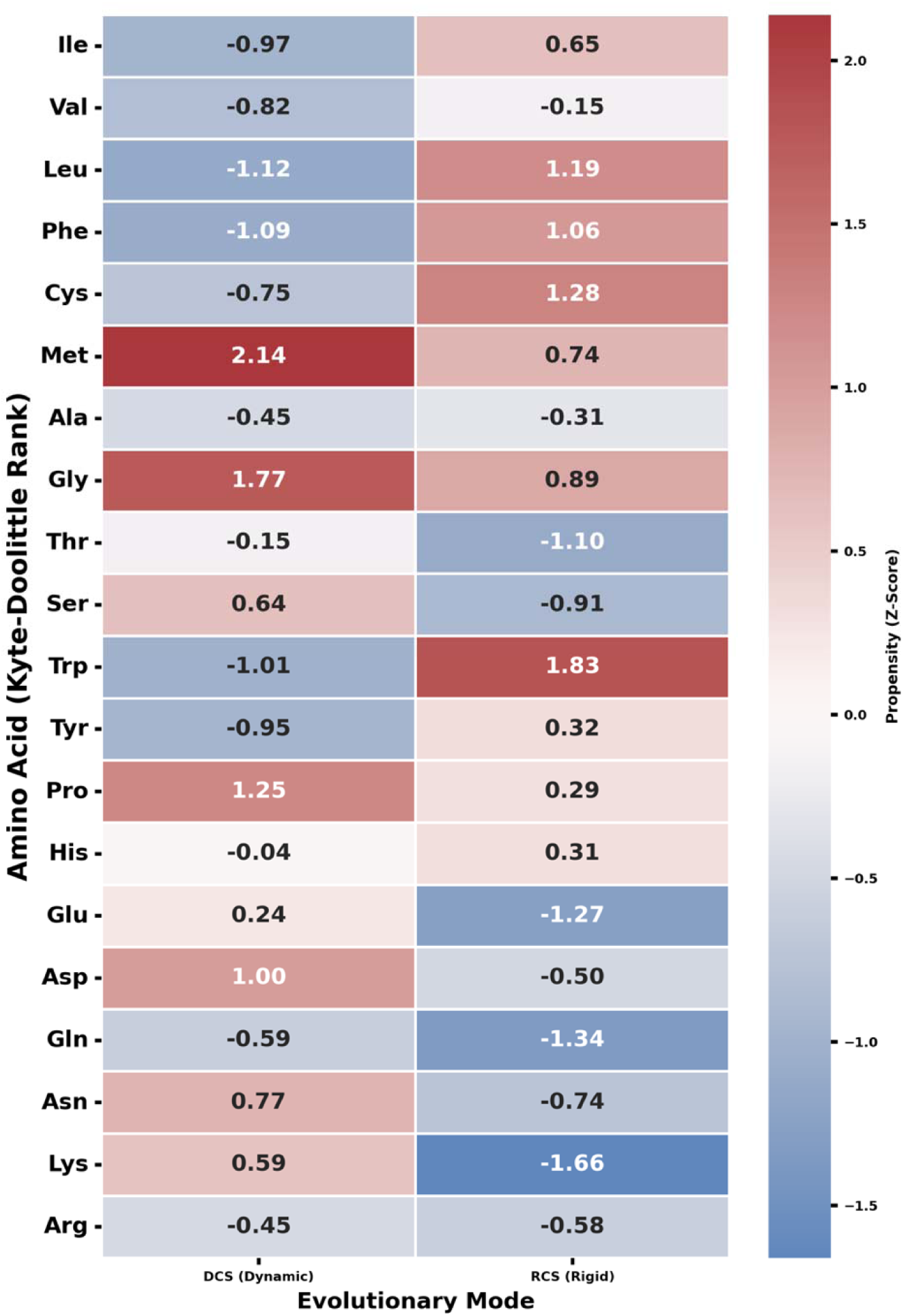
Amino Acid Propensity Heatmap. The heatmap is visualizing the enrichment of each residue type within high-scoring deciles of DCS and RCS, calculated from the pan-proteome dataset.

As expected, this chemical map contrasts greatly with the Rigid Conserved Score (RCS) profile (Figure 2, right column). The RCS metric specifically isolates the structural core -mainly hydrophobic, showing extreme enrichment for Tryptophan (Z=1.83), Cysteine (Z=1.28), and Leucine (Z=1.19). By resolving these distinct chemical codes, we provide a molecular basis for the divergent pathogenic profiles observed: the loss of a Tryptophan or Cysteine leads to global unfolding (high RCS), whereas the loss of a conserved Glycine or Proline leads to the arrest of functional motion (high DCS). Interestingly, our recent investigations into the *QTY-Code*, a design strategy replacing hydrophobic residues with polar ones [8], have independently highlighted this fundamental separation between chemical packing and functional kinematics. Evolutionary profiling [9] and variant analyses of water-soluble QTY-variants [10, 11] reveal that the protein core often tolerates radical chemical substitutions provided the volumetric shape is preserved, distinguishing these sites from those governed by strict kinematic necessity. Furthermore, comparative molecular dynamics and AlphaFold [13, 25] modeling of water-soluble variants [26, 27, 28] demonstrate that essential fluctuation profiles and ligand-binding interactions persist even when the hydrophobic core is chemically erased. Complementary to our earlier findings, the hydrophobic residues replaced in the QTY-code protocol (Leucine, Valine, Isoleucine, and Phenylalanine) [8] display a great difference: they are strongly depleted in the DCS while showing greater scores in the RCS (Figure 2, top 4 row). These findings reinforce our observation that the evolutionary constraints preserving the protein’s folded state are physically distinct from, and operate independently of, the topological constraints encoding its essential motions.

### Determinants of Mutational Intolerance

Consistent across both datasets, structural rigidity emerged as the primary driver of mutational intolerance. In correlation (Supplementary Figures 5 and 6) and <u>R</u>eceiver <u>O</u>perating <u>C</u>haracteristic (ROC) analysis [29] (Figure 3d), the Rigid Conserved Score achieved state-of-the-art performance in discriminating pathogenic variants compared to AlphaMissense, yielding an <u>A</u>rea <u>U</u>nder the <u>C</u>urve (AUC) of 0.75. This performance is comparable to raw conservation (AUC=0.74) but significantly outperforms the Dynamic Conserved Score (AUC=0.51). Given the inherent imbalance in clinical datasets, we further evaluated their practical utility using Top-K Precision analysis (Figure 3e). At the top 1% of scores, RCS achieves a precision of 0.80 for identifying lethal variants (AlphaMissense > 0.90), significantly surpassing the 0.61 precision achieved by raw conservation. As shown in the heatmap of (Figure 2, right column), residues with high RCS are strictly enriched in hydrophobic amino acids (Tryptophan Z=1.83, Leucine Z=1.19) and stabilizing disulfides (Cysteine Z=1.28). This chemical signature likely confirms that RCS specifically isolates the lethal structural core, where mutations are likely to trigger global misfolding and loss of function.

**Figure 3.**
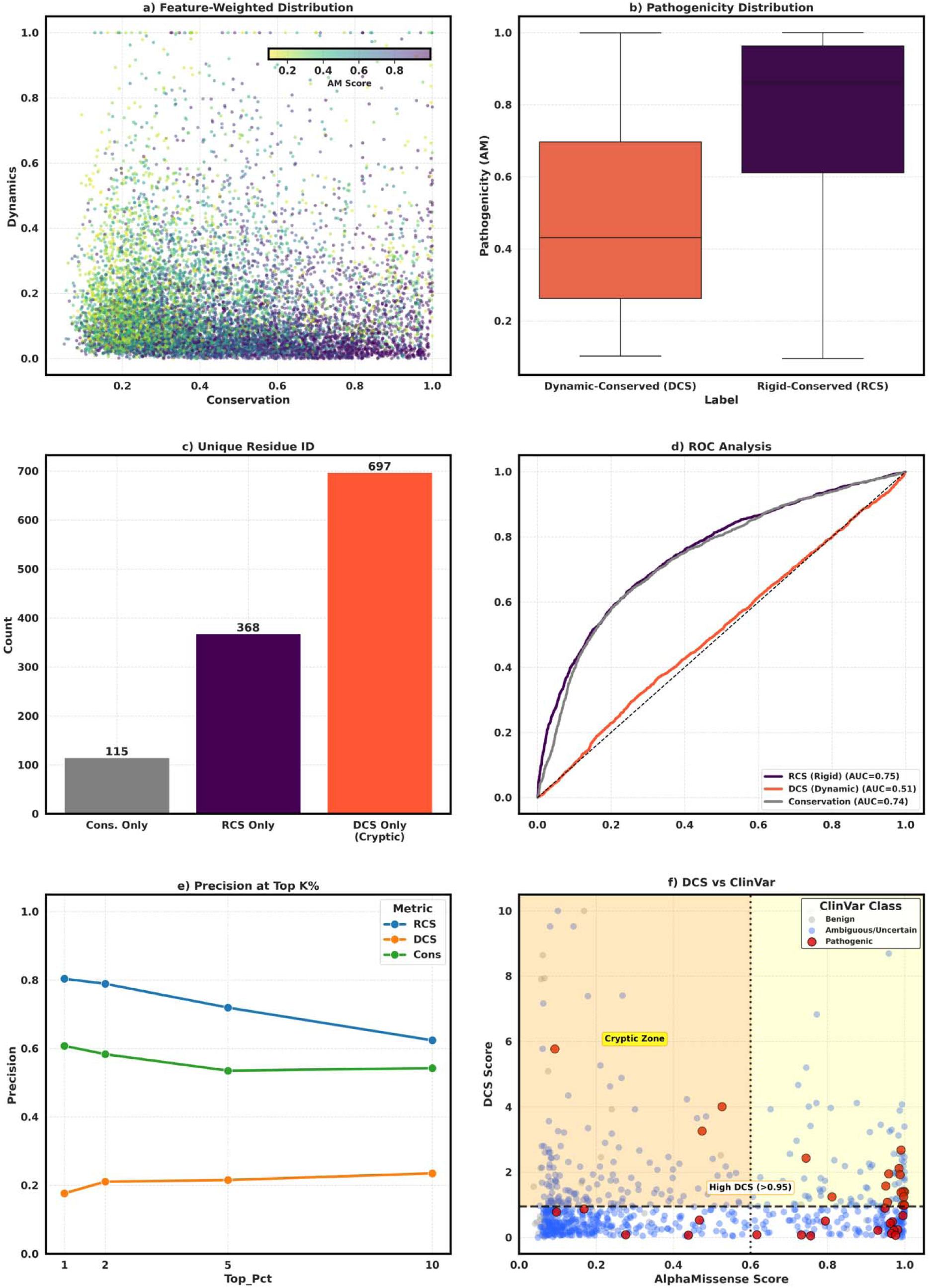
Statistical Analysis of Dynamics-aware Evolutionary Profiling Metrics. **(a)** Feature-weighted landscape of AlphaMissense scores plotted against evolutionary conservation and structural dynamics. High pathogenicity scores (purple) cluster predominantly in regions of high conservation and low dynamics. **(b)** Distribution of AlphaMissense pathogenicity scores across biophysical classes. **(c)** Unique residue identification within the top 10% of scores. DCS identifies the largest distinct set of residues (n=697), sites missed by conservation and rigidity metrics alone. **(d)** <u>R</u>eceiver <u>O</u>perating <u>C</u>haracteristic (ROC) analysis benchmarking RCS, DCS, and Conservation against AlphaMissense. **(e)** Precision at Top-K% analysis, at the top 1% of scores, RCS achieves a precision of 0.80 for identifying AlphaMissense Pathogenic variants (AM > 0.90). **(f)** Clinical validation of DCS against human disease variants. Scatter plot mapping 737 ClinVar variants onto the dynamics-pathogenicity landscape. Variants are colored by clinical classification: Pathogenic (red), Benign (gray), and Ambiguous (blue). The DCS Discovery Zone (shaded orange; DCS > 0.95, AlphaMissense < 0.6) isolates variants located in flexible functional regions that are misclassified as benign or ambiguous by AlphaMissense predictors.

On the other hand, a fundamental signal inversion was observed when examining the relationship between protein dynamics and pathogenicity, visualized in the feature-weighted landscape of (Figure 3a). Protein flexibility (raw dynamics) correlates negatively with pathogenicity (Spearman ρ≈-0.38, Figure 1b), statistically validating the hypothesis that flexible residues generally tolerate mutation. However, crucially, when dynamics is contextually weighted by conservation via the DCS metric, this correlation shifts significantly toward neutrality (ρ≈0.08). This represents a statistical shift relative to raw dynamics, demonstrating that DCS may filter out the benign background noise of random loops. Unlike general-flexibility, which is protective (negative correlation), the conserved-flexibility captured by DCS is functionally constrained. This inversion marks a biophysical transition from mutational tolerance to mutational sensitivity (in dynamic sites).

### The Population of Soft-Conserved Sites

In our analysis of the human-proteome, DCS isolated a distinct population of 697 unique residues in the top decile that were completely missed by standard conservation analysis (Figure 3c). This metric shows a subset of high-scoring candidates identified exclusively by our dynamics-aware method and not by baseline conservation. This subset represents a critical blind spot in current genomic profiling: residues that are evolutionarily maintained not for their packing density, but for their specific role in the dynamic ensemble. While the Rigid Conserved Score serves as a potent predictor of structural stability, the Dynamic Conserved Score identifies a cryptic, orthogonal class of sites that remains largely invisible to traditional conservation metrics

In addition, these Dynamic-Conserved sites exhibit a pathogenic profile (from AlphaMissense [14] scores) that is distinct, as visualized in the boxplot distributions of (Figure 3b). While the Rigid-Conserved core is associated with high-confidence pathogenicity, the sites identified uniquely by DCS occupy an intermediate pathogenic regime (Figure 3b). This intermediate profile suggests that mutations at these sites are mostly not catastrophic. Unlike core mutations which likely trigger global unfolding and degradation, mutations in these dynamic regions likely impair specific *soft* functions, such as allosteric regulation, signal transduction, or binding kinetics, without abolishing the protein fold entirely [5]. This finding can redefine the concept of druggability in the context of genetic variation. High-RCS sites are often *hard* targets where modulation is difficult. In contrast, the *soft* pharmacophores identified by DCS represent tunable nodes within the protein network. By highlighting these mechanically essential sites, our framework could provide a filter for identifying allosteric pockets and regulatory switches that are amenable to subtle pharmacological intervention.

### The Cluster of Dynamic Constraint (Biophysical Goldilocks Zone)

We employed unsupervised K-Means clustering [30] for defining the biophysical architecture of the proteome. This algorithm partitions the multidimensional data into discrete phase spaces based on variance minimization, effectively mapping the proteome into distinct biophysical regimes. This analysis revealed a unique Dynamic-Conserved cluster, characterized by the paradoxical combination of high evolutionary conservation and intrinsic structural flexibility. Statistical validation via the Mann-Whitney U test [31] confirms that this cluster represents an intermediate pathogenic zone. Visualized as the green belt in the radial biophysical map (Figure 4), residues in this zone are significantly more pathogenic (AlphaMissense scores) than the evolutionary buffer region (dense non-conserved, cyan) with a highly significant p-value (p≈1.58×10-11). As illustrated in (Figure 4), while the Rigid-Conserved residues (magenta) cluster near the high-constraint center, the Dynamic-Conserved residues (yellow) occupy an intermediate orbit. This position signifies a regime of balanced constraint, physically distinct from the outer periphery of high biophysical freedom where mutational tolerance is elevated. This distinction is critical because current static predictors heavily weight structural packing [3]. In the absence of dynamic data, a flexible conserved residue often appears chemically ambiguous or structurally disordered. We term this the “Biophysical Goldilocks Zone” (Figure 4) by analogy to the astronomical habitable zone, a region that is capable of supporting life because it is neither too hot nor too cold. In our proteomic landscape, this corresponds to an intermediate regime that is neither strictly rigid nor non-conserved, but maintains the precise balance of flexibility and constraint required for function. By delineating this intermediate phase space, our Dynamics-Aware Profiling highlights a population of residues that are functionally brittle, possessing the flexibility required for motion yet lacking the evolutionary redundancy of the buffer zone. Identifying this cluster is essential for resolving Variants of Uncertain Significance (VUS), as it flags variants that appear structurally benign but are mechanically lethal.

**Figure 4.**
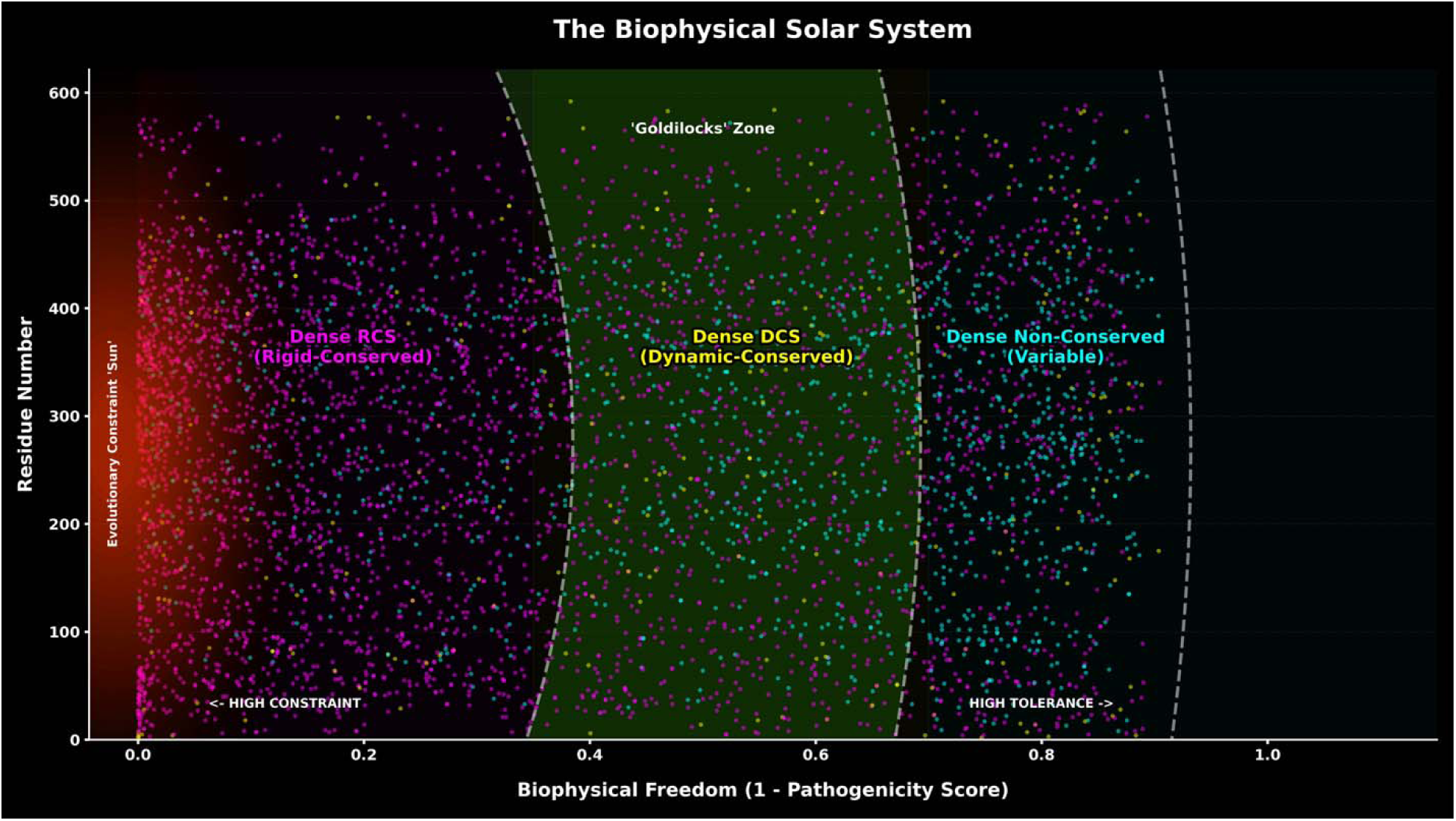
Conceptual Mapping of Pathogenicity Landscapes. The relationship between dynamics-aware evolutionary scores and pathogenicity (AlphaMissense) is visualized using a gravitational analogy. Here, distance from the center correlates with mutational tolerance (1 - Pathogenicity) (Biophysical Freedom). <u>R</u>igid-<u>C</u>onserved <u>S</u>core (RCS) residues (Magenta) form a high-constraint core, analogous to a gravitational center with lethal intolerance to mutation. <u>D</u>ynamic-<u>C</u>onserved <u>S</u>core (DCS) residues (Yellow) inhabit an intermediate *conceptual* approximate biophysical Goldilocks Zone (Green) of balanced constraint (from Figure 3b). The periphery (Cyan) characterized by low conservation and high tolerance for genetic variation. The background scatter points represent individual residues from the human proteome dataset (Magenta=High RCS, Yellow=High DCS, Cyan=Variable)

### RCS and DCS as Complementary Filters for Variant Interpretation

To benchmark the performance of our metrics against state-of-the-art static predictors, we evaluated the comparative practical utility of RCS and DCS versus AlphaMissense [14] using Top-K Precision analysis [32] (Figure 3e). This metric simulates a real-world prioritization scenario by asking: “If a scientist investigates only the top K percent of highest-scoring variants from each model, what fraction are true positives?”. In our analysis (Figure 3e), RCS achieved a precision of 0.80 within the top 1% of scores, significantly surpassing the 0.61 precision achieved by raw conservation alone (Gray line). This substantial enhancement confirms that RCS effectively strips away the noise of non-structural conservation, providing a high-confidence filter for structural variants where mutations are in rigid sites. Conversely, Dynamic Conserved Score serves as a specialized filter for a more subtle class of variants in dynamic sites. While DCS targets a smaller population (precision ≈0.18 at top 1% due to the rarity of these sites) (Figure 3e), its value lies in specificity rather than broad sensitivity. By identifying sites that are evolutionarily constrained yet structurally mobile, DCS shows variants that appear benign in static crystal structures but are critical in the dynamic ensemble. This complementarity allows for an additional classification of genomic risk: High RCS variants can be flagged as likely destabilizers, while high DCS variants can be prioritized as likely disruptors of allosteric signaling or mechanism. This dual-channel approach moves beyond the monolithic view of pathogenicity, offering a granular framework for interpreting variants on their specific biomechanical impact.

Because current predictors inherently award stability [1, 2, 3], they may frequently classify dynamics regions as variable or benign. These essential dynamic pharmacophores are discarded during the initial stages of target prioritization. By highlighting these cryptic pockets, our framework significantly expands the druggable proteome. Traditional drug discovery has largely focused on the rigid active site, which offers high affinity but often poor selectivity. However, the soft sites identified by DCS likely represent the tunable nodes of the allosteric network. These sites are ideal targets for the development of allosteric modulators (which can fine-tune protein function) without the toxicity associated with complete structural inhibition. The identification of these unique targets provides new maps (Figure 5) for targeting the unknown area of the proteome.

**Figure 5.**
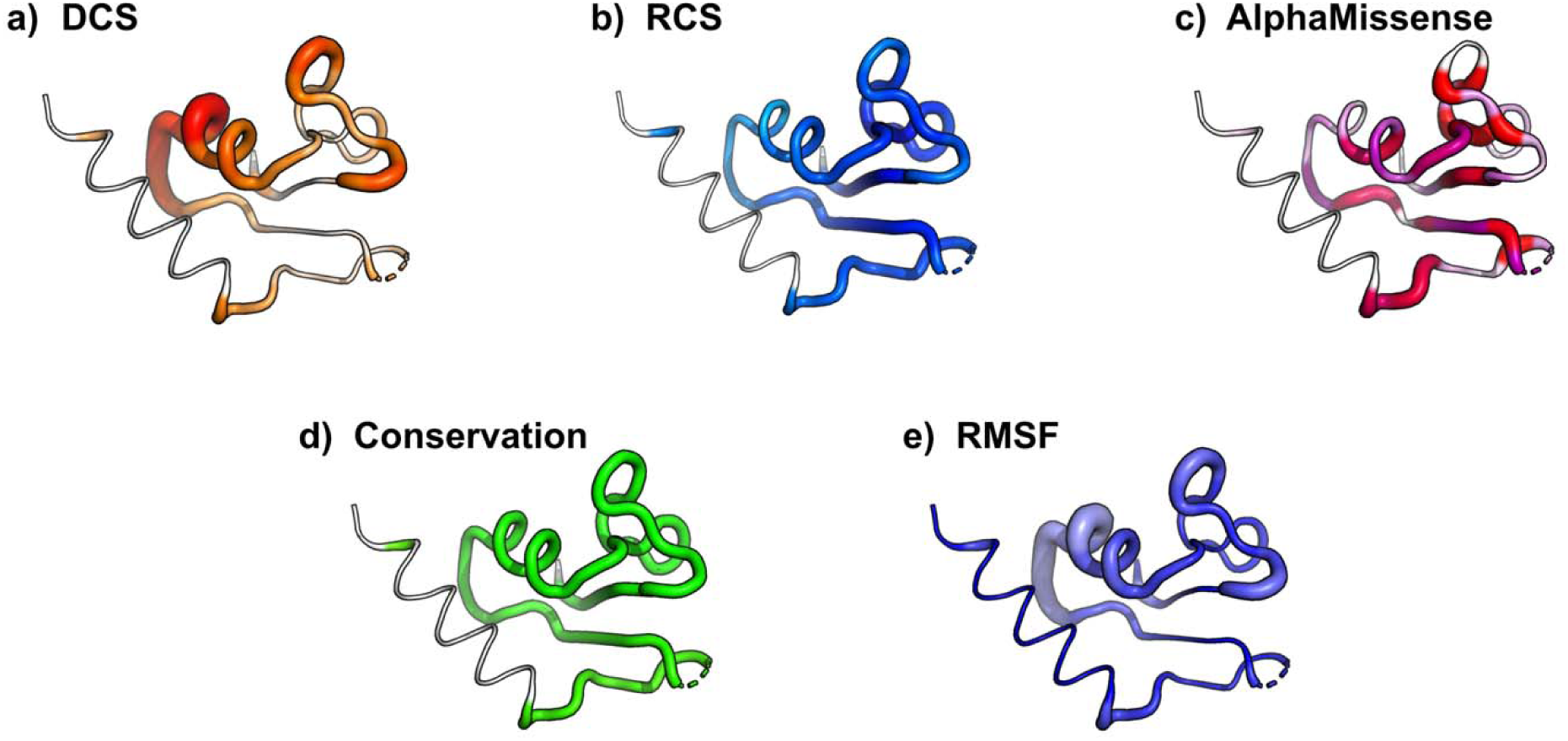
3D structural mapping of a representative human protein. Structural maps of human asparaginyl-tRNA synthetase (PDB ID: 4zya chainA), illustrating how DCS identifies specific flexible sites that are distinct from the rigid scaffold but potentially critical for allosteric function. **(a)** DCS mapped to tube thickness (putty representation). **(b)** Equivalent mapping for RCS (Blue=High RCS), **(c)** Comparison with AlphaMissense pathogenicity predictions, **(d)** raw conservation from EVcouplings, and **(e)** averaged RMSF values from ATLAS database.

Furthermore, this biological adaptability extends beyond internal mechanics to environmental sensing, a dimension entirely missing in static structural studies. Our recent investigations into pH-dependent protein dynamics [33, 34] demonstrate that evolution often uses specific flexible residues to modulate protein behavior in response to electrostatic shifts. Because computational methods typically rely on structures solved at a single fixed pH, they systematically fail to detect these environmentally sensitive residues. For instance, in Junctional Adhesion Molecules, we identified strictly conserved residues that may act as pH sensors by altering their fluctuation profiles, a functional role invisible to standard stability metrics [33]. Similarly, in Synaptogyrins, we showed that nearly identical structural backbones can encode vastly different dynamic responses to membrane acidity [34]. By capturing the evolutionary constraint on motion rather than just static position, DCS can successfully flag these environmental sensors, offering a novel strategy for designing therapeutics that selectively target tissues with specific pH profiles, such as the acidic microenvironments of tumors.

### Clinical Validation on Human Disease Variants

To validate the utility of our biophysical metrics in a clinical setting, we mapped 737 human variants from the ClinVar database [18] onto our computed landscapes. This analysis confirmed that DCS captures a pathogenic mechanism distinct from static structural stability. As visualized in (Figure 5f), the Dynamic Conserved Score (DCS) demonstrated a unique capacity to identify pathogenic variants located in flexible functional regions, a class of mutations often misclassified as benign by structural predictors like AlphaMissense.

We defined *DCS Discovery Zone* (DCS > 0.95, AlphaMissense < 0.6, ClinVar Pathogenic/Likely Pathogenic) to isolate these cryptic pathogenic events where the evolutionary constraint is dynamic rather than structural, but classified as non-pathogenic in AlphaMissense. Our analysis revealed specific examples such as PGK1p.Lys30Thr (DCS=3.26) and NARS1 p.Lys60Thr (DCS=4.01), which are clinically pathogenic but received ambiguous or benign scores from AlphaMissense (0.47 and 0.53, respectively) (Table 1).. These findings highlight the DCS metric as a critical filter for resolving Variants of Uncertain Significance (VUS) in flexible regions, where disease is driven by the arrest of molecular mechanics rather than thermodynamic instability.

**Table 1.** High Threshold Cryptic Pathogenic Discoveries within ClinVar from DCS (> 0.95) and RCS (> 6.0) metrics, compared to AlphaMissense (< 0.6).

| Protein | Variant | DCS (0-10)<sup>1</sup> | RCS (0-10)<sup>2</sup> | Alpha Missense<sup>3</sup> | ClinVar<sup>4</sup> | Cryptic Metric<sup>5</sup> |
| --- | --- | --- | --- | --- | --- | --- |
| NARS1 | p.Thr17Met | 5.78 | 7.96 | 0.09 (benign) | Likely pathogenic | DCS and RCS |
| NARS1 | p.Lys60Thr | 4.01 | 7.82 | 0.53 (uncertain) | Likely pathogenic | DCS and RCS |
| PGK1 | p.Lys30Thr | 3.26 | 0.62 | 0.47 (uncertain) | Likely pathogenic | DCS |
| SPRTN | p.Tyr117Cys | 0.53 | 6.18 | 0.47 (uncertain) | Pathogenic | RCS |

<sup>1</sup>Dynamic Conserved Score (DCS) is defined as the product of conservation and normalized dynamics.
<sup>2</sup>Rigid Conserved Score (RCS) is defined as the product of conservation and the inverse of normalized dynamics.
<sup>3</sup>AlphaMissense score of the variant (0-10).
<sup>4</sup>ClinVar Classification.
<sup>5</sup>DCS (> 0.95) and/or RCS (> 6.0) metrics, compared to AlphaMissense (< 0.6).

Complementing this analysis, the Rigid Conserved Score (RCS) served as a control for lethal structural defects. Consistent with its role in core stability, RCS identified a population of pathogenic variants in the RCS Discovery Zone (RCS > 6.0, AlphaMissense < 0.6). Notably, variants such as SPRTN p.Tyr117Cys (RCS=6.18) and NARS1 p.Thr17Met (RCS=7.96) scored highly on our rigid-conservation scale despite receiving low or ambiguous pathogenicity scores from AlphaMissense (0.47 and 0.09, respectively) (Table 1) (Supplementary Table 2). This suggests that while DCS captures the dynamic sites, RCS rescues the structural variants that are underestimated by sequence-based methods alone.

### Co-evolution as a Mutational Buffer

Finally, we investigated the role of higher-order network effects by defining the <u>D</u>ynamics <u>Co</u>-evolutionary Cou<u>p</u>ling <u>S</u>core (DCopS), calculated as the product of evolutionary enrichment (co-evolutionary coupling [20]) and normalized dynamics. Unsupervised Principal Component Analysis (PCA) of this coupled landscape (Supplementary Figure 7) reveals that Co-evolutionary Coupling (PC1, 58.3% variance) and Dynamics (PC2, 41.7% variance) form orthogonal axes of constraint. This independence is further confirmed by the correlation matrix in (Supplementary Figure 7b), which shows that DCopS operates distinctly from the rigid scaffold, exhibiting a negative correlation with the <u>R</u>igid <u>Co</u>-evolutionary Cou<u>p</u>ling <u>S</u>core (RCopS) (ρ=-0.26). We assessed the relationship between this score and pathogenicity, observing a negative correlation with the coupling component (ρ≈-0.13).

Unlike the isolated dynamic sites identified by DCS, residues with high dynamic coupling appear to act as mutational buffers. This is quantitatively validated by the coreelation and <u>R</u>eceiver <u>O</u>perating <u>C</u>haracteristic (ROC) analysis (Supplementary Figures 8, 9, 10), where DCopS yields an AUC of 0.42. Biophysical profiling (Supplementary Figure 11) provides the molecular basis for this buffering capacity. The DCopS aminoacid signature has a unique enrichment of Tryptophan (Z=2.10), Glutamine (Z=1.14), Serine (Z=1.10), and Lysine (Z=1.05). As expected, this chemical profile contrasts with the rigid core (depleted in Cysteine, Z=-1.06), suggesting these regions are not disulfide-locked scaffolds but rather promiscuous interaction interfaces (Trp) or flexible polar loops (Ser, Gln, Lys) (Supplementary Figure 11) These regions possess the thermodynamic capacity to absorb sequence variation through compensatory structural shifts across the coupled network. DCopS identifies 346 unique residues that are distinct from those flagged by rigidity (RCopS) or pure coupling (Supplementary Figure 8). By distinguishing the buffer of the network from the machinery of the protein, DCopS may prevent the misclassification of flexible, highly coupled regions as pathogenic.

Furthermore, in our recent investigations into Glutamate Transporters (EAATs) [35] and <u>V</u>esicular <u>M</u>ono<u>a</u>mine <u>T</u>ransporters (VMATs) [36], we identified truncation splice isoforms that precisely mimic the highly coupled oligomerization interfaces of their canonical counterparts via molecular dynamics. Interestingly, these isoforms leverage evolutionarily highly coupled binding hotspots to compete with full-length monomers, thereby acting as molecular buffers that regulate transporter assembly [35, 36]. Just as DCopS residues may absorb mutational stress to preserve the network, these isoform interfaces absorb the dynamic drive for oligomerization. This suggests that coupled dynamic networks are versatile evolutionary modules, capable of being repurposed via alternative splicing.

### Future Scopes and the Potential Applications

The mathematical orthogonality of the Rigid- and Dynamic-Conserved Scores establishes a powerful dual-filter strategy for precision medicine, that resolve a long-standing ambiguity in variant interpretation. The identification of dynamic-conserved residues via the Dynamics-aware Evolutionary Profiling fundamentally expands the boundaries of the druggable proteome, rendering the cryptic allosteric network visible to rational design. Traditional pharmacological strategies have predominantly targeted high RCS sites, the rigid highly conserved active sites or ATP-binding pockets, which are structurally ubiquitous across families and prone to off-target toxicity. However, our analysis reveals that high DCS regions represent a distinct class of *soft* pharmacophores: tunable nodes that are mechanically essential but dynamically distinct from the rigid sites. Because these sites function dynamically rather than static scaffolds, they offer ideal targets for the development of allosteric modulators rather than structural inhibitors. By targeting these potential flexible regulatory switches, future therapeutic projects can effectively manipulate specific protein motions or force inactive conformations without competing with endogenous ligands. This strategy is particularly promising for targets previously considered undruggable, such as transcription factors, where the active site is too shallow yet the allosteric regulation is mediated by the specific dynamic networks.

Beyond therapeutic intervention, the orthogonality of stability and dynamics addresses a critical bottleneck in precision medicine, the interpretation of <u>V</u>ariants of Uncertain <u>S</u>ignificance (VUS). Current clinical classifiers may flag variants in flexible regions as benign due to the assumption that structural disorder implies functional dispensability. Our results demonstrate that this is a false negative, residues of conserved flexibility can be pathogenic, but their mechanism of action differs fundamentally. By stratifying variants into rigid (high RCS) and dynamic (high DCS) classes, clinicians can better predict patient phenotypes. Implementing this dual-channel scoring system into molecular boards could significantly improve the resolution of genetic diagnoses and guide the selection of conformation-specific therapies.

Using a cohort of 737 ClinVar [18] variants, we confirmed that our dual-score system potentially resolves the biophysical ambiguity of genetic disease. We show that while high RCS residues mark the *hard* structural scaffold rigid residues, the high DCS residues highlight the *soft* allosteric dynamic residues. The identification of clinically pathogenic variants with high DCS but low AlphaMissense scores, such as NARS1 p.Thr17Met, validates the existence of a dynamic pathogenic regime that must be integrated into future genomic interpretation pipelines. To facilitate broad community adoption and ensure reproducibility, we have developed an open-access web server, *ADEPT* (<u>A</u>utomated <u>D</u>ynamics-aware <u>E</u>volutionary <u>P</u>rofiling <u>T</u>ool) (https://www.karagolresearch.com/adept) that automates the calculation of these scores from standard input data (Supplementary Figure 12). Our results implies that successful *de novo* protein design can optimize a multi-objective function, ensuring folding while tuning DCS to engineer specific permissible motions. Future iterations of generative protein models could incorporate these constraints to design synthetic enzymes with pre-programmed flexibility profiles/proteins that are not only stable but also possess the requisite breathing motions for high-efficiency catalysis, moving beyond the static optimization of the folded state to the dynamic optimization of the proteins.

## Methods

### Dataset Curation and Filtering Protocols

The construction of our dataset was driven by the requirement for high-fidelity structural and dynamic data. We established two distinct cohorts to validate our framework across different biological contexts. The primary *Cross-Species Alpha-Helical* cohort consists of 93 membrane and globular proteins derived from the ATLAS Molecular Dynamics database (https://www.dsimb.inserm.fr/ATLAS) [19] and RCSB Protein Data Bank (https://www.rcsb.org/) [37]. To ensure that the subsequent evolutionary profiling mapped to stable molecular dynamics data and similar protein class, we applied alpha-helical filtering criteria: a Root Mean Square Deviation (RMSD) threshold of < 2Å during equilibration, a minimum chain length of 150 residues, and a secondary structure composition of 40–70% alpha-helicity [38]. The second *Homo Sapiens Proteome* cohort was curated to assess clinical relevance, comprising 58 medium-length human proteins (400-600 residues) identified by cross-referencing ATLAS entries with the UniProt database [39].

### Molecular Dynamics Trajectory Analysis

For each protein in the curated dataset, we extracted per-residue <u>R</u>oot <u>M</u>ean <u>S</u>quare <u>F</u>luctuation (RMSF) profiles from three independent simulation replicas from ATLAS database [19]. These replicas were averaged to minimize stochastic sampling noise and capture the canonical flexibility profile of the backbone. Because raw RMSF values are inherently scale-dependent and influenced by protein size and packing density, we applied Min-Max normalization to the averaged RMSF vectors. This transformation scales the dynamic profile of every protein to a standardized 0-1 range (Norm_Dynamics), allowing for the direct statistical comparison of flexibility patterns across diverse protein architectures without bias from global structural variance.

### Evolutionary and Co-Evolutionary Profiling

To map the evolutionary landscape, we generated deep evolutionary profiles using the EVcouplings framework (https://v2.evcouplings.org) [20]. <u>M</u>ultiple <u>S</u>equence <u>A</u>lignments (MSAs) were constructed using the Jackhmmer algorithm [40] against the UniProt database, employing iterative search parameters to maximize sequence diversity while minimizing gaps. Site-specific conservation scores were calculated based on the Shannon entropy [41] of each alignment column. To capture higher-order evolutionary constraints, we have used co-evolutionary coupling scores (Enrichment) [20], which quantify the statistical dependency between residue pairs. These coupling metrics serve as a proxy for functional interactions, identifying residues that may be individually variable but are constrained by the requirements of the residue interaction network.

### Formulation of Dynamics-Aware Metrics

We developed a suite of orthogonal metrics to mathematically uncouple the biophysical forces of stability and motion (results). The <u>R</u>igid <u>C</u>onserved <u>S</u>core (RCS) is defined as the product of conservation and the inverse of normalized dynamics (1-Norm_Dynamics),. Conversely, the <u>D</u>ynamic <u>C</u>onserved <u>S</u>core (DCS) is defined as the product of conservation and normalized dynamics, isolating dynamic pharmacophores by selecting for residues that are strictly conserved yet structurally mobile. To interrogate the role of network effects, we defined the <u>D</u>ynamic <u>Co</u>-evolutionary Cou<u>p</u>ling <u>S</u>core (DCopS) and <u>R</u>igid <u>Co</u>-evolutionary Cou<u>p</u>ling <u>S</u>core (RCopS) as the product of evolutionary coupling enrichment with normalized dynamics and inverse dynamics, respectively. All resulting scores were Min-Max scaled to a 0-10 range per protein to standardize the dynamic range for downstream statistical applications.

### Analysis Infrastructure

To handle the substantial computational load required for analysis, all computational pipelines were executed within the Google Colab Pro Plus high-performance environment. The workflow was accelerated using TPUv2-8 Tensor Processing Units, supported by 334.6 GB of RAM and 225.3 GB of system storage. This infrastructure facilitated high-throughput matrix operations and the rapid processing of large-scale evolutionary data. Custom Python and R scripts [42] utilizing the scikit-learn [43], pandas [44], and scipy [45] libraries were developed to execute the statistical analyses and generate high-dimensional visualizations. The scripts are openly available for result re-creation. PyMOL is used for 3D structural visualization and mapping [46]. For Figure 5, the structure of human asparaginyl-tRNA synthetase (PDB ID: 4ZYA [37, 47], chain A) was used.

### Statistical Analysis

To validate the hypothesis that stability and dynamics represent independent evolutionary channels, we performed dimensionality reduction on the six-dimensional feature space. <u>P</u>rincipal <u>C</u>omponent <u>A</u>nalysis (PCA) [21] was employed to project the proteomic data onto orthogonal axes, this relationship was further confirmed using Spearman rank correlation [48] coefficients, which assess monotonic relationships between variables. Our analysis specifically tested the independence of RCS and DCS to ensure that the Dynamic-Conserved signal was not merely a residual artifact of the Rigid-Conserved signal.

Our framework was also benchmarked against the AlphaMissense database [14], a state-of-the-art deep learning predictor of pathogenicity, scores accessed through HageLab’s AlphaMissense webpage (https://alphamissense.hegelab.org) [49]. We classified residues into categories, Pathogenic(score > 0.90) and Benign(score < 0.20), discarding ambiguous predictions to ensure dataset score purity. We utilized <u>R</u>eceiver <u>O</u>perating <u>C</u>haracteristic (ROC) [28] analysis to calculate the global discrimination performance. Complementarily, we assessed the metrics via Top-K Precision analysis [32]. This technique calculates the precision of the model at specific high-confidence thresholds (top 1%, 2%, 5%, and 10% of scores), simulating a real-world drug discovery scenario where resources are limited to validating only the highest-ranking targets.

To define the biophysical states of the proteome independent of clinical labels, we applied K-Means clustering [30] to the global dataset, using normalized conservation and dynamics as input features. The statistical significance of the differences in pathogenicity between our clusters was assessed using the Kruskal-Wallis H test followed [22] by Dunn’s post-hoc test [50]. Additionally, the chemical logic of these clusters was interrogated by calculating the propensity of specific amino acids to populate each cluster with Mann-Whitney U [31] tests.

### Clinical Validation

To validate the metrics against clinical observation, we mapped human variants from the ClinVar database (https://www.ncbi.nlm.nih.gov/clinvar/) [18] onto our computed landscapes. This cohort included single nucleotide missense variants with established clinical significance (Pathogenic/Likely Pathogenic, Benign/Likely Benign, or Uncertain Significance) mapped to the 58 human proteins in our dataset. We integrated pathogenicity predictions by assigning AlphaMissense scores to each variant based on the specific amino acid substitution. We defined discovery zones to identify cryptic pathogenic variants missed by static prediction: the DCS Discovery Zone (DCS > 0.95, AlphaMissense < 0.6, ClinVar Pathogenic/Likely Pathogenic) and the RCS Discovery Zone (RCS > 6.0, AlphaMissense < 0.6, ClinVar Pathogenic/Likely Pathogenic). The enrichment of pathogenic variants in these zones versus the background was assessed using descriptive statistics.

### Web Server Implementation

To ensure the reproducibility of our findings and facilitate broad community adoption, we developed *ADEPT* (<u>A</u>utomated <u>D</u>ynamics-aware <u>E</u>volutionary <u>P</u>rofiling <u>T</u>ool) (https://www.karagolresearch.com/adept), an open-access web server, with client-side processing, similar to our previous Evolutionary Statistics Toolkit [51]. The application is architected as a secure client-side tool to maintain data privacy. It accepts standard CSV inputs containing per-residue RMSF and evolutionary conservation/coupling data and automatically computes the full suite of dynamics-aware scores (DCS, RCS, DCopS, RCopS). The server integrates the 3Dmol.js library [52] to provide interactive, browser-based visualization.

## Supplementary information (SI)

Supplementary_Information.pdf

## Data Availability

The molecular dynamics simulation results database, with corresponding PDB files accessible via the ATLAS database (https://www.dsimb.inserm.fr/ATLAS) and RCSB Protein Data Bank (https://www.rcsb.org/). Conservation scores for these proteins are available at the EvCouplings server (https://v2.evcouplings.org), while individual AlphaMissense scores can be found at HageLab’s AlphaMissense webpage (https://alphamissense.hegelab.org). ClinVar variant data is available at (https://www.ncbi.nlm.nih.gov/clinvar/). Researchers can use our open-access tool *ADEPT* (https://www.karagolresearch.com/adept) to create scores from MD and evolutionary data. A comprehensive list of proteins analyzed in this study, along with Python and R scripts used for data preparation and statistical analysis, as well as additional statistical data, can be accessed at https://github.com/karagol-taner/Dynamics-aware-Evolutionary-Profiling

## Author Contributions

Both authors (T.K., and A.K.) equally contributed to this manuscript. Conceptualization, T.K., A.K.; methodology, T.K., A.K.; software, T.K., A.K., AlphaMissense, EvCouplings, ATLAS database, R, Python; data curation, T.K., A.K.; validation, T.K., A.K.; formal analysis, T.K., A.K.; investigation, T.K., A.K., writing - original draft preparation, T.K., A.K., visualization, T.K., A.K.

## Additional Statements

Request for additional information should be addressed to corresponding authors, Taner Karagöl, or Alper Karagöl.

