## Supplementary_Information.pdf for "Dynamics-aware Evolutionary Profiling Uncouples Structural Rigidity from Functional Motion to Enable Enhanced Variant Interpretation"

### Table of Contents

1. Table S1. List of analyzed Homo sapiens and cross-species proteins. More detailed information is available in the data repository. Pg. 2-4
2. Figure S1. Top-ranked cross-species alpha-helical proteins based on highest average DCS, derived from available molecular dynamics data and structural analysis (see Methods). Pg. 5
3. Figure S2. Top-ranked cross-species alpha-helical proteins based on highest average RCS, derived from available molecular dynamics data and structural analysis (see Methods). Pg. 6
4. Figure S3. Top-ranked human medium-size proteins based on highest average DCS, derived from available molecular dynamics data and structural analysis (see Methods). Pg. 7
5. Figure S4. Top-ranked human medium-size proteins based on highest average RCS, derived from available molecular dynamics data and structural analysis (see Methods). Pg. 8
6. Figure S5. Comparison between Rigid Conserved Score (RCS) and AlphaMissense scores. Pg. 9
7. Figure S6. Comparison between Dynamic Conserved Score (DCS) and AlphaMissense scores. Pg. 10
8. Figure S7. Descriptive Statistics of DCopS and RCopS. Pg. 11
9. Figure S8. Statistical Analysis of Dynamics-aware Evolutionary Coupling Profiling Metrics. Pg. 12
10. Figure S9. Comparison between Rigid Co-evolutionary Coupling Score (RCopS) and AlphaMissense scores. Pg. 13
11. Figure S10. Comparison between Dynamics Co-evolutionary Coupling Score (DCopS) and AlphaMissense scores. Pg. 14
12. Figure S11. Amino Acid Propensity Heatmap of DCopS and RCopS metrics. Pg. 15
13. Figure S12. Screenshot of ADEPT (Automated Dynamics-aware Evolutionary Profiling Tool) web page. Pg. 16
14. Table S2. List of available variants from our analysis that are also available in the ClinVar database (likely pathogenic and/or pathogenic variants only). Pg. 17-18

**Table S1.** List of analyzed *Homo sapiens* and *cross-species* proteins. More detailed information is available in the data repository. (https://github.com/karagol-taner/Dynamics-aware-Evolutionary-Profiling)

| ID[1] | PDB ID[2] | UniProt ID[3] |
|---|---|---|
| 1 | 2xolB | O30077 |
| 2 | 3fhfA | Q58134 |
| 3 | 3bjdB | Q9HY91 |
| 4 | 3letB | Q87P32 |
| 5 | 2pfxA | Q1GDR9 |
| 6 | 4xb4A | P74102 |
| 7 | 3oyvA | A7M120 |
| 8 | 2o70F | A1L259 |
| 9 | 3nr1A | Q8N4P3 |
| 10 | 2imsA | Q16611 |
| 11 | 3rznA | Q9NZD2 |
| 12 | 3fcnA | Q2RPE2 |
| 13 | 2o74F | A1L259 |
| 14 | 2zcaB | Q53VY0 |
| 15 | 5w4aB | - |
| 16 | 2p54A | Q07869 |
| 17 | 3jz9A | Q5ZSQ3 |
| 18 | 1j77A | Q9RGD9 |
| 19 | 4hfvA | Q5ZUE7 |
| 20 | 5w4dB | - |
| 21 | 1r8mE | Q99418 |
| 22 | 5cofA | Q1R1X2 |
| 23 | 2fozA | Q9NX46 |
| 24 | 1c1kA | P13342 |
| 25 | 1ah7A | P09598 |
| 26 | 2bbrA | Q98325 |
| 27 | 3nkeB | Q46896 |
| 28 | 5fiaB | Q5ZXU7 |
| 29 | 2incA | O87798 |
| 30 | 4qn8A | Q5ZRR7 |
| 31 | 3p5pA | Q41594 |
| 32 | 1z1lA | O00408 |
| 33 | 3jzaB | Q9H0U4 |
| 34 | 3m7dA | Q977W1 |
| 35 | 2fefB | Q9I1R6 |
| 36 | 2osaA | P15882 |
| 37 | 1fczA | P13631 |
| 38 | 2x96A | Q10714 |
| 39 | 4uc8A | P03418 |
| 40 | 2fefA | Q9I1R6 |
| 41 | 4qn8B | Q5ZRR7 |
| 42 | 3hfwA | P54922 |
| 43 | 2gnoA | Q9WZM9 |
| 44 | 2y1qA | P37571 |
| 45 | 1wbeA | P68265 |
| 46 | 5fbfA | P24021 |
| 47 | 3lyfD | D3K5I7 |
| 48 | 1vdkB | P84127 |
| 49 | 7bqiA | Q9BQS8 |
| 50 | 3eqxA | Q8E9K5 |
| 51 | 2ovjA | Q9H0H5 |
| 52 | 2ixmA | Q15257 |

| | | |
|---|---|---|
| 53 | 1iomA | Q5SIM6 |
| 54 | 1w2wA | Q06489 |
| 55 | 1d2tA | Q9S1A6 |
| 56 | 4z9pA | P18272 |
| 57 | 3owtA | P11938 |
| 58 | 2zd2B | A1B060 |
| 59 | 3lgbB | P20457 |
| 60 | 4ny3A | Q15257 |
| 61 | 5b1qB | Q9WT50 |
| 62 | 2dwkA | Q9D394 |
| 63 | 3m6zA | Q977W1 |
| 64 | 5x1uB | Q5WZ95 |
| 65 | 6exaB | Q5ZYC7 |
| 66 | 5b1qA | Q9WT50 |
| 67 | 3owtB | P11938 |
| 68 | 1w2wE | Q06489 |
| 69 | 5zmoA | Q9L0M9 |
| 70 | 4n67A | A0A0H3LV04 |
| 71 | 1qgiA | P33673 |
| 72 | 7e2sA | P72583 |
| 73 | 5lc8A | Q8RNT4 |
| 74 | 1y6iA | P72583 |
| 75 | 4ervA | Q15413 |
| 76 | 2abkA | P0AB83 |
| 77 | 5muxB | P45859 |
| 78 | 1s9uA | Q8ZPK0 |
| 79 | 5j47A | P03072 |
| 80 | 5muxC | P45859 |
| 81 | 1tezB | P05327 |
| 82 | 1hbnE | P11558 |
| 83 | 7aqxA | P26332 |
| 84 | 1e6yD | P07962 |
| 85 | 8b3wA | M4T0T6 |
| 86 | 3p3oA | Q70KH6 |
| 87 | 4djaA | A9CH39 |
| 88 | 6mm7C | P05132 |
| 89 | 3cmlA | Q6UDW7 |
| 90 | 5imuA | O05446 |
| 91 | 3bqkA | Q6UDW7 |
| 92 | 8cxlA | A7KH32 |
| 93 | 2wsiA | P38913 |
| h1 | 1by2A | Q08380 |
| h2 | 1dtdB | P48052 |
| h3 | 1ef1C | P26038 |
| h4 | 1ef1D | P26038 |
| h5 | 1egwB | Q02078 |
| h6 | 1egwC | Q02078 |
| h7 | 1ewfA | P17213 |
| h8 | 1f5nA | P32455 |
| h9 | 1fczA | P13631 |
| h10 | 1fs1A | Q13309 |
| h11 | 1fs1C | Q13309 |
| h12 | 1lj5A | P05121 |
| h13 | 1m9zA | P37173 |
| h14 | 1mlwA | P17752 |
| h15 | 1nkpD | P01106 |

| | | |
|---|---|---|
| h16 | 1o4kA | P12931 |
| h17 | 1wojA | P09543 |
| h18 | 2cb5A | Q13867 |
| h19 | 2f60K | O00330 |
| h20 | 2osaA | P15882 |
| h21 | 2p54A | Q07869 |
| h22 | 2pq8A | Q9H7Z6 |
| h23 | 2vkpB | Q96KE9 |
| h24 | 2wyaA | P54868 |
| h25 | 2wzbA | P00558 |
| h26 | 2wzoA | Q3YBR2 |
| h27 | 2xdgA | Q02643 |
| h28 | 2xvsA | Q8N0Z6 |
| h29 | 2znrA | Q96FJ0 |
| h30 | 3bqpB | P07602 |
| h31 | 3lblA | Q00987 |
| h32 | 3luiB | Q15036 |
| h33 | 3m66A | Q96E29 |
| h34 | 4bt9A | P13674 |
| h35 | 4bt9B | P13674 |
| h36 | 4ddpA | Q14457 |
| h37 | 4fqnB | Q9BSQ5 |
| h38 | 4gv2A | Q9Y6F1 |
| h39 | 4hkdB | Q13188 |
| h40 | 4jnuA | Q7Z3B4 |
| h41 | 4jnuB | Q7Z3B4 |
| h42 | 4jo7A | Q9BVL2 |
| h43 | 4jo7C | Q9BVL2 |
| h44 | 4xglA | O15091 |
| h45 | 4ykdA | Q9BSQ5 |
| h46 | 4zyaA | O43776 |
| h47 | 4zyaB | O43776 |
| h48 | 5opmA | P49902 |
| h49 | 5v03B | Q9H2S1 |
| h50 | 5w2fA | P41214 |
| h51 | 5wbxB | Q9H2S1 |
| h52 | 6mdwA | Q9H040 |
| h53 | 6mdxA | Q9H040 |
| h54 | 6mi3B | Q9Y6K9 |
| h55 | 6uxeB | Q9Y697 |
| h56 | 6y43A | Q8NI99 |
| h57 | 7jflC | Q14653 |
| h58 | 7jflD | Q14653 |

[1]Protein IDs 1–93 are cross-species alpha-helical proteins; IDs h1–h58 are Homo sapiens medium-sized proteins. Proteins were selected from available molecular dynamics trajectories within the ATLAS database (https://www.dsimb.inserm.fr/ATLAS) (methods).
[2]PDB ID from the RCSB PDB (https://www.rcsb.org/)
[3]UniProt ID of the protein.

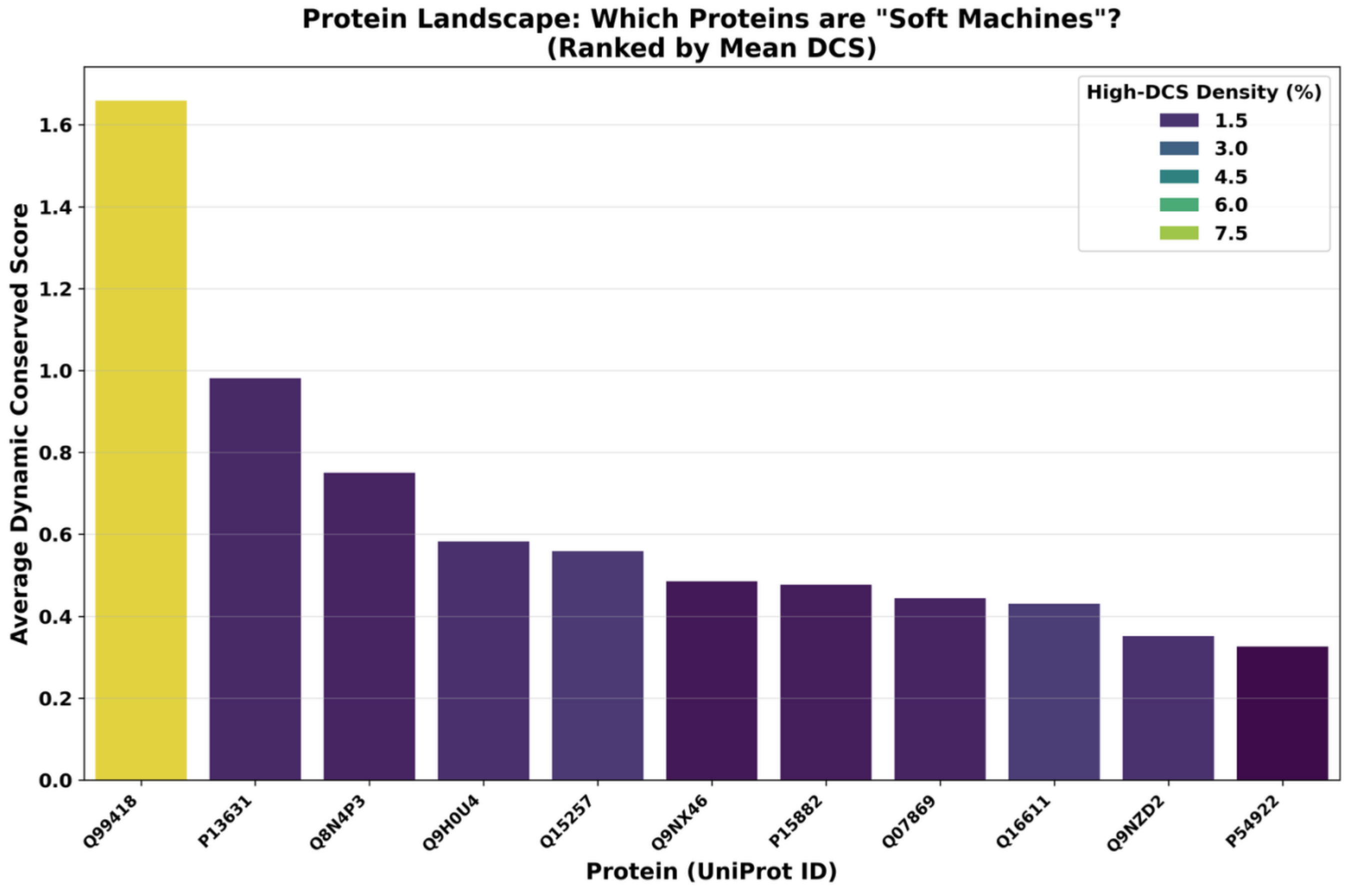


**Figure S1.** Top-ranked cross-species alpha-helical proteins based on highest average DCS, derived from available molecular dynamics data and structural analysis (see Methods).

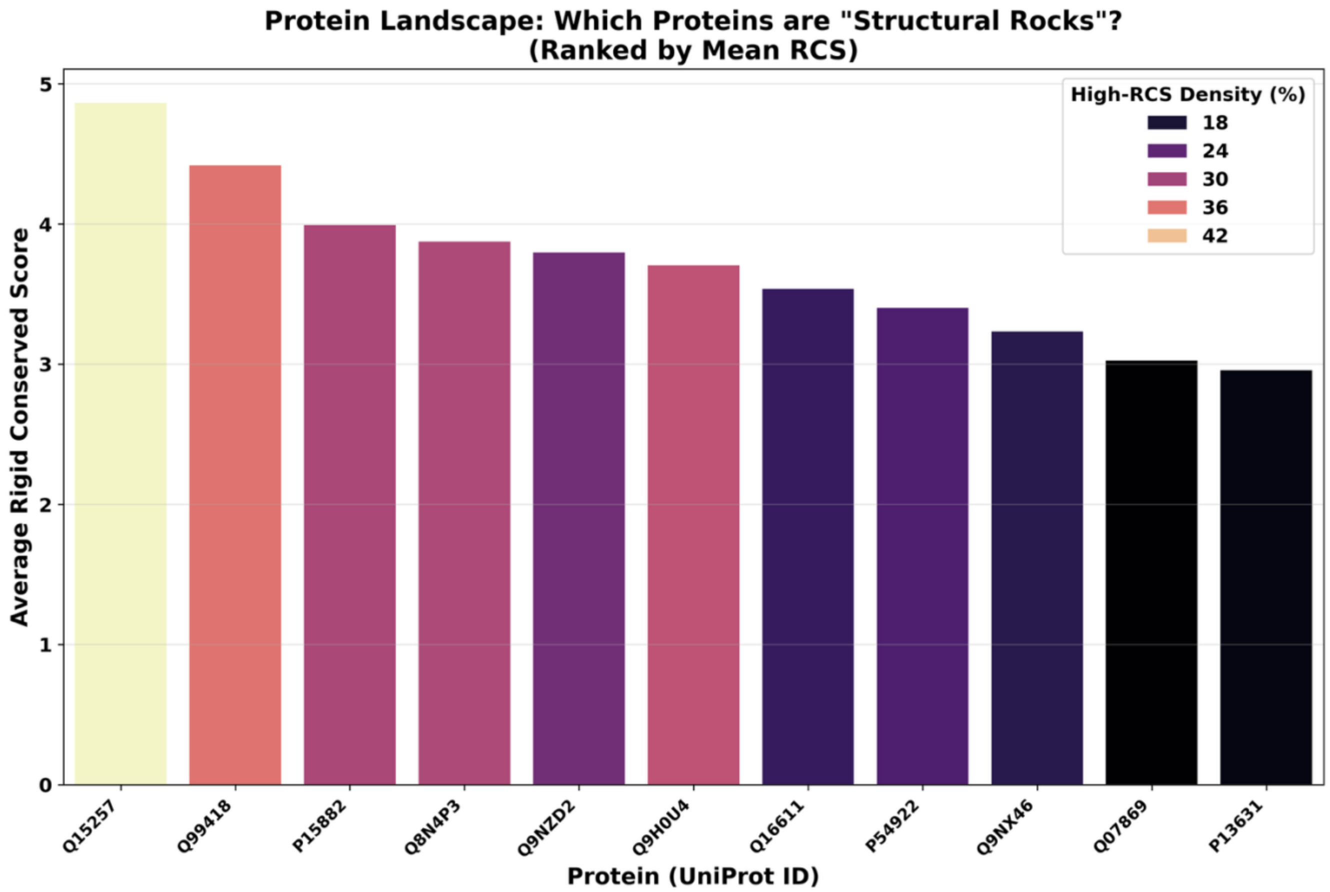


**Figure S2.** Top-ranked cross-species alpha-helical proteins based on highest average RCS, derived from available molecular dynamics data and structural analysis (see Methods).

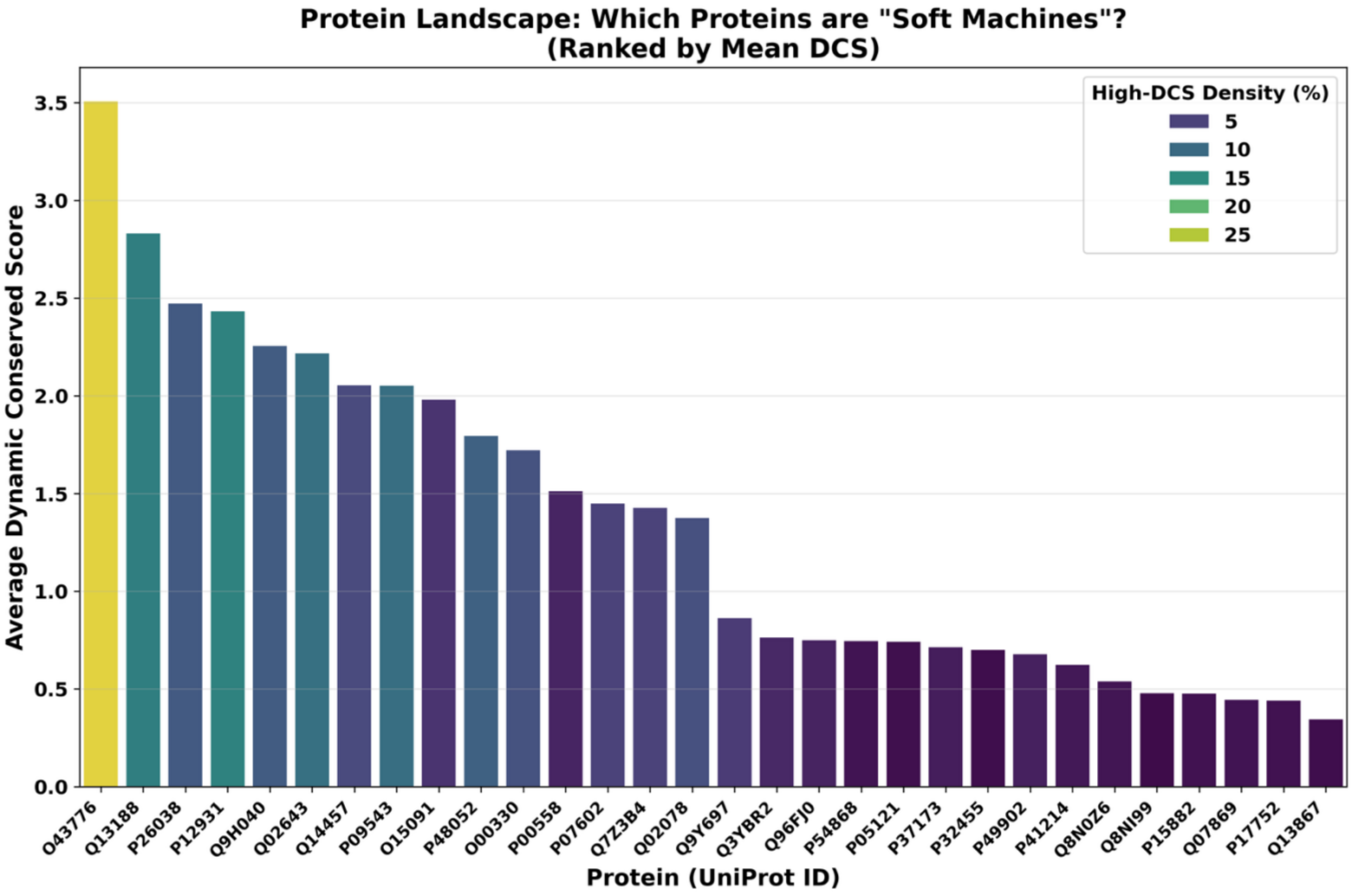


**Figure S3.** Top-ranked human medium-size proteins based on highest average DCS, derived from available molecular dynamics data and structural analysis (see Methods).

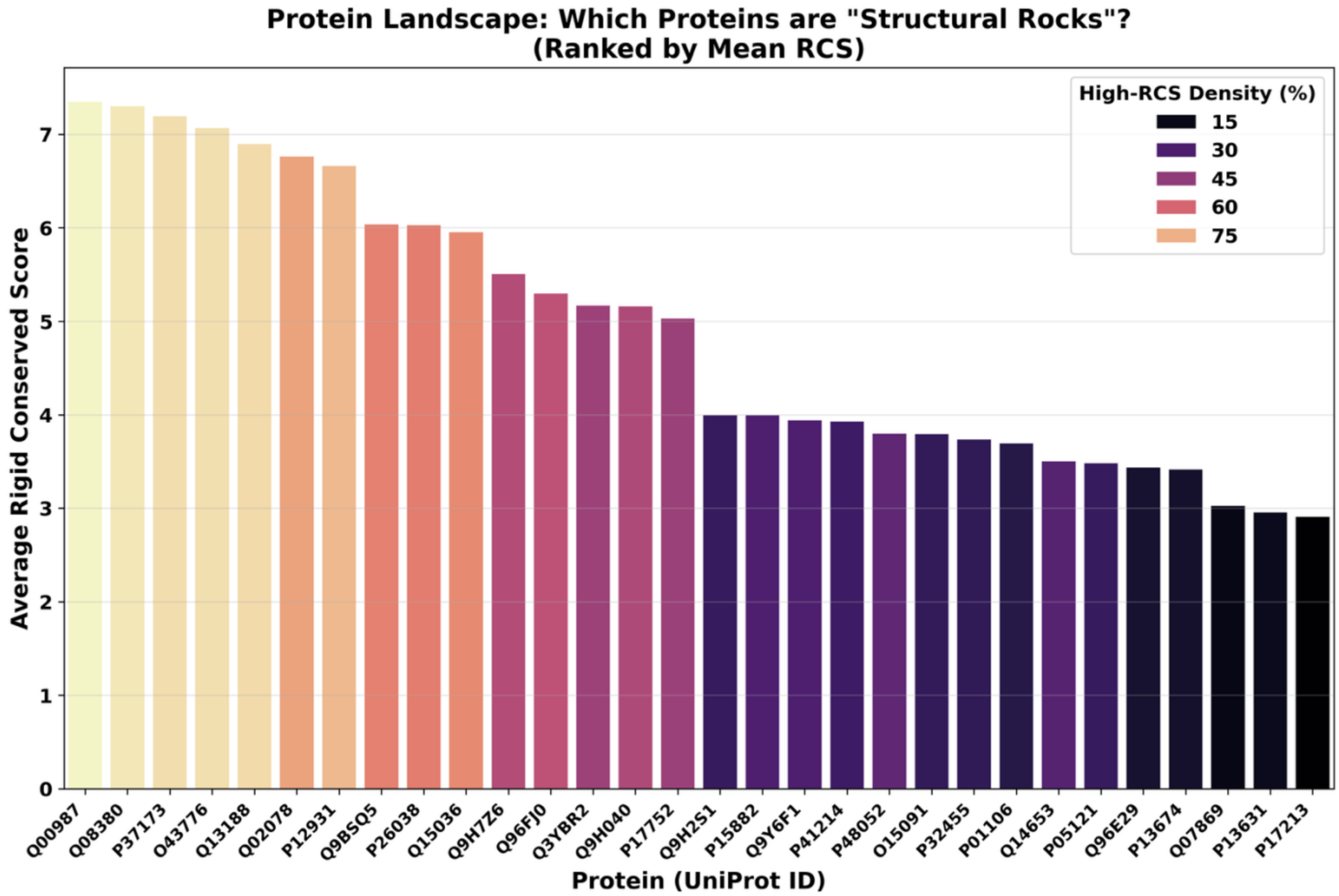


**Figure S4.** Top-ranked human medium-size proteins based on highest average RCS, derived from available molecular dynamics data and structural analysis (see Methods).

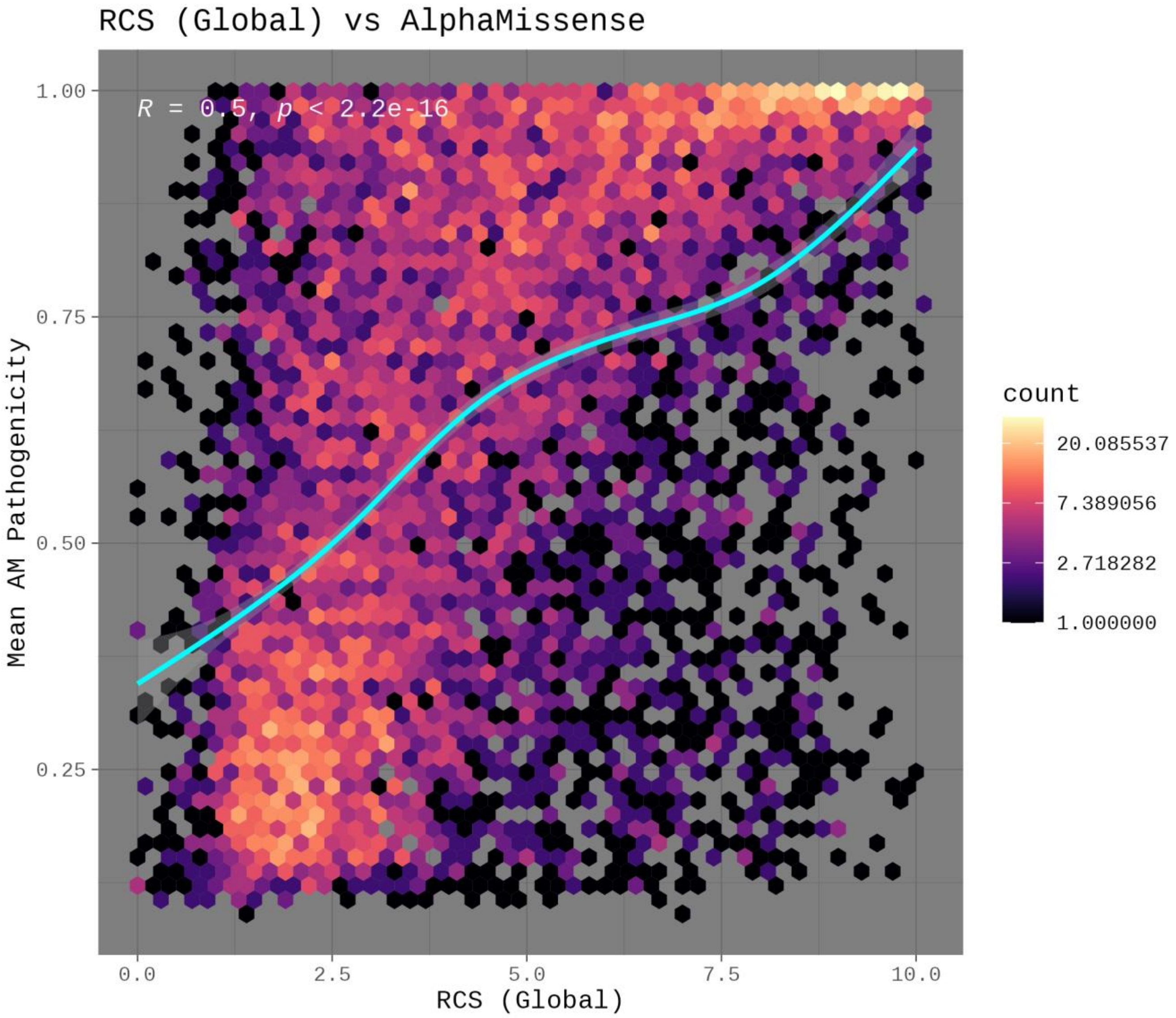


**Figure S5.** Comparison between Rigid Conserved Score (RCS) and AlphaMissense scores.

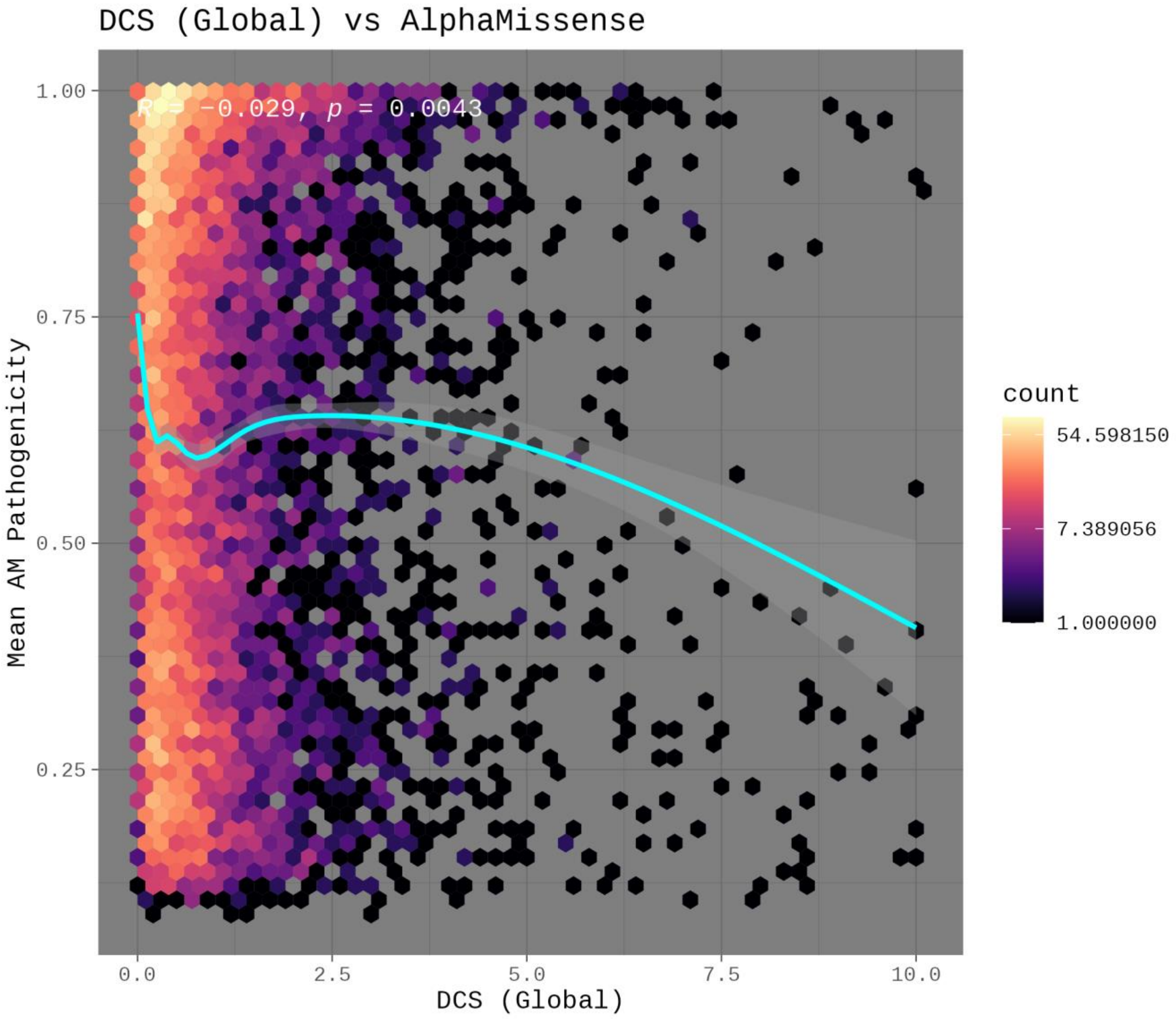


**Figure S6.** Comparison between Dynamic Conserved Score (DCS) and AlphaMissense scores.

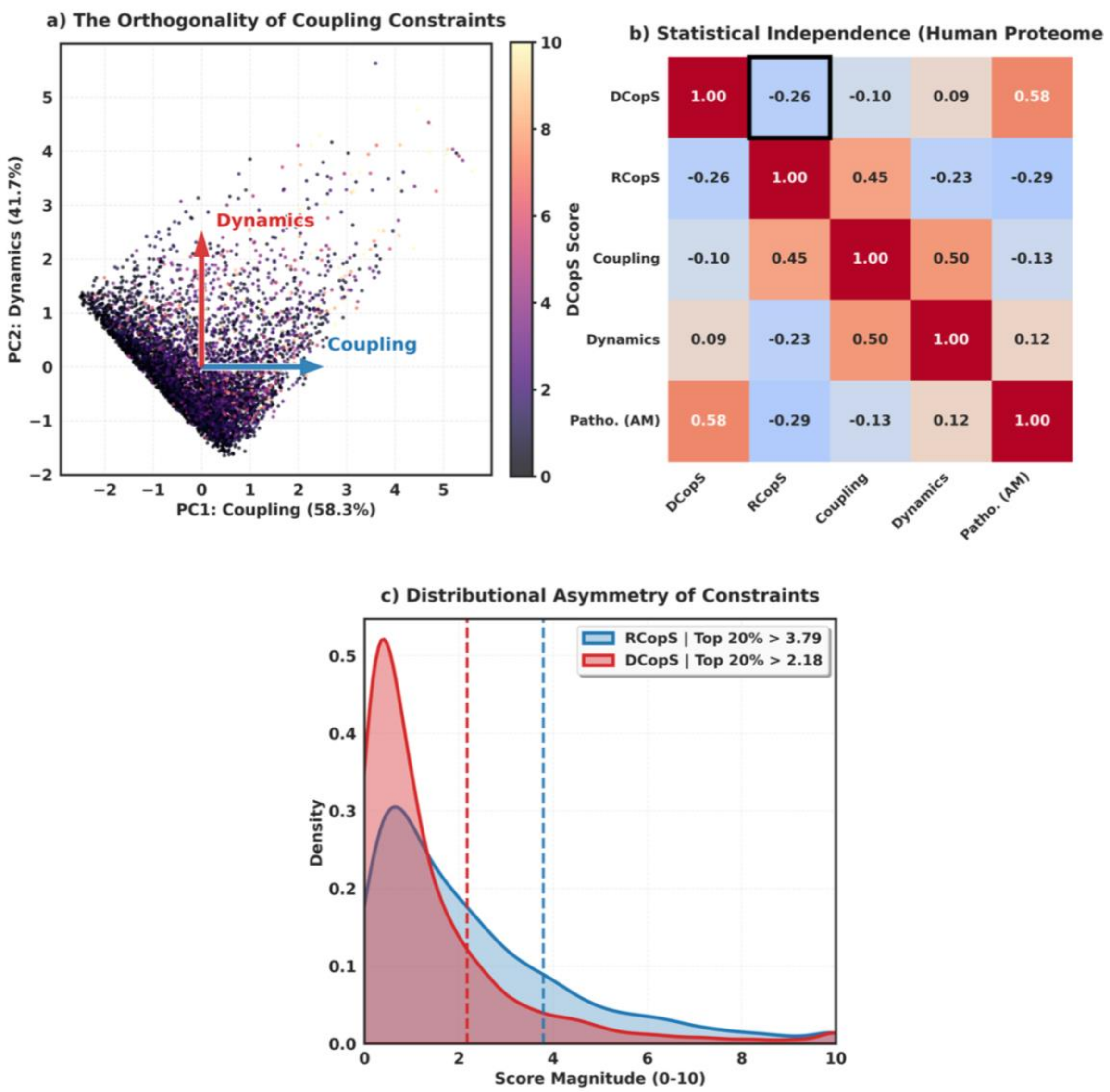


**Figure S7.** Descriptive Statistics of DCopS and RCopS**.**
(a) Principal Component Analysis (PCA) biplot of the pan-proteome residue landscape (93 alpha-helical proteins), illustrating the relationship between Evolutionary Coupling and Structural Dynamics. b) Spearman correlation matrix quantifying the relationships between derived metrics. (c) Density plots of DCopS and RCopS scores.

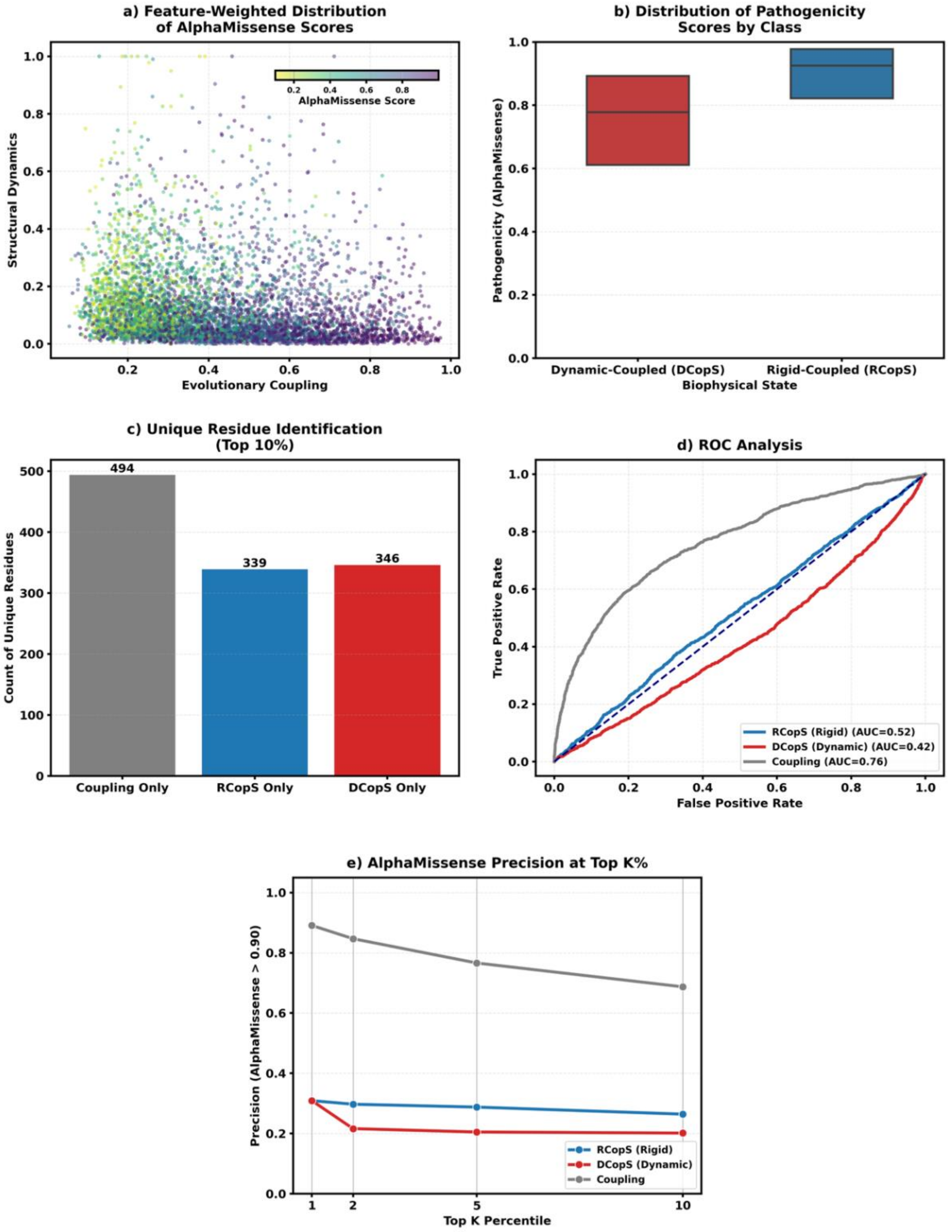


**Figure S8.** Statistical Analysis of Dynamics-aware Evolutionary Coupling Profiling Metrics. (a) Feature-weighted landscape of AlphaMissense scores plotted against evolutionary coopling and structural dynamics. (b) Distribution of AlphaMissense pathogenicity scores across biophysical classes. (c) Unique residue identification within the top 10% of scores. (d) Receiver Operating Characteristic (ROC) analysis (e) Precision at Top-K% analysis.

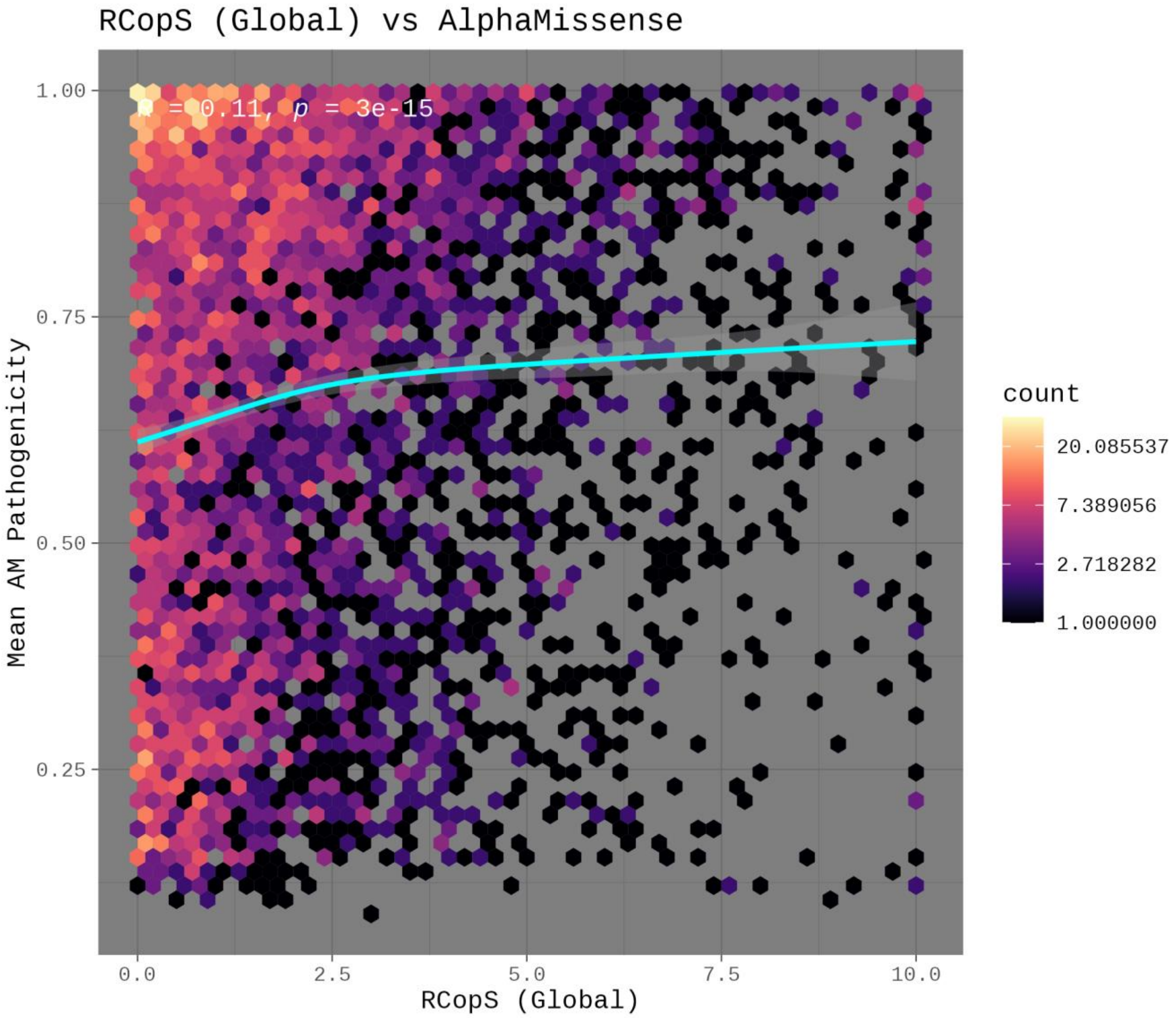


**Figure S9.** Comparison between Rigid Co-evolutionary Coupling Score (RCopS) and AlphaMissense scores.

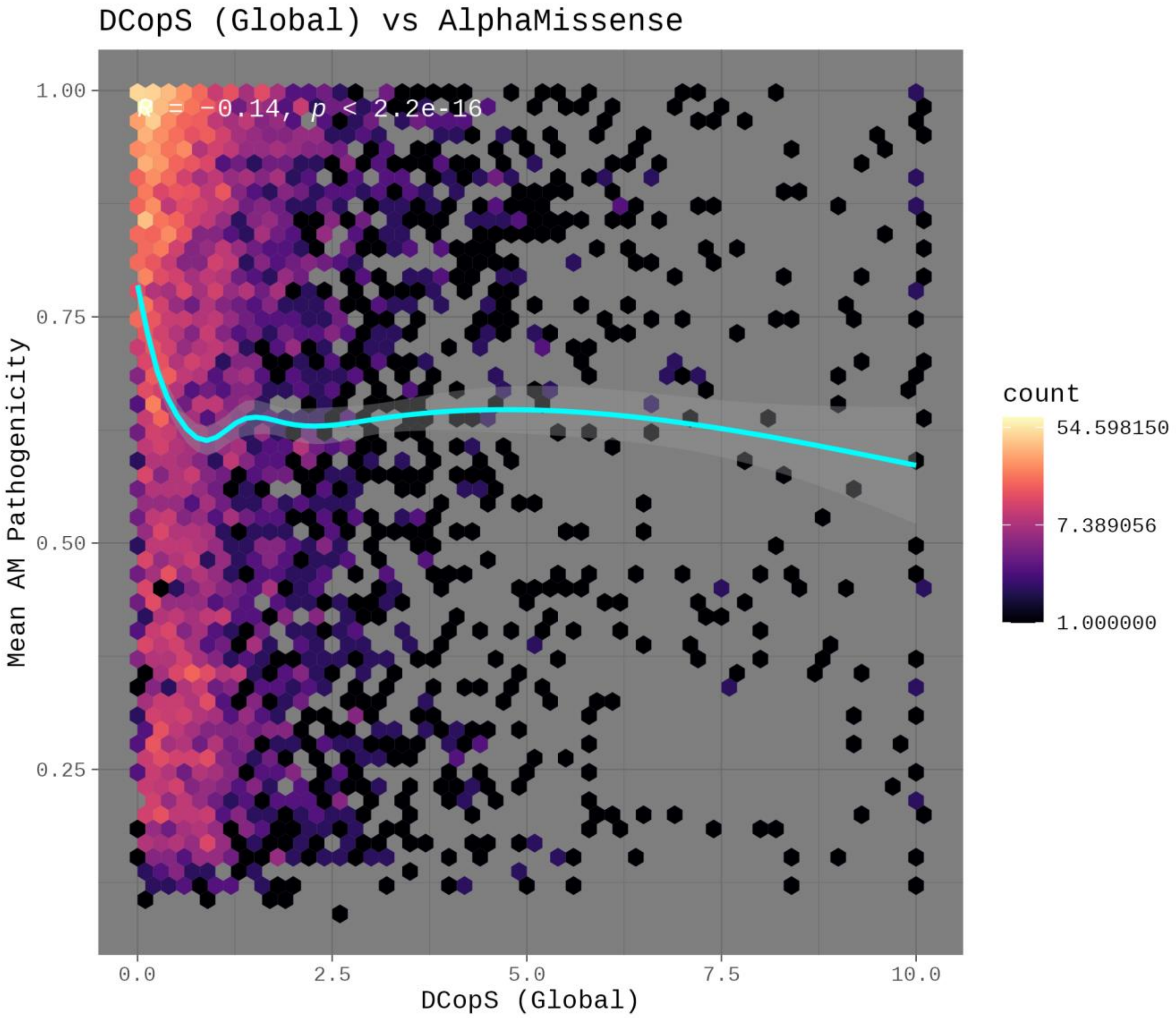


**Figure S10.** Comparison between Dynamics Co-evolutionary Coupling Score (DCopS) and AlphaMissense scores.

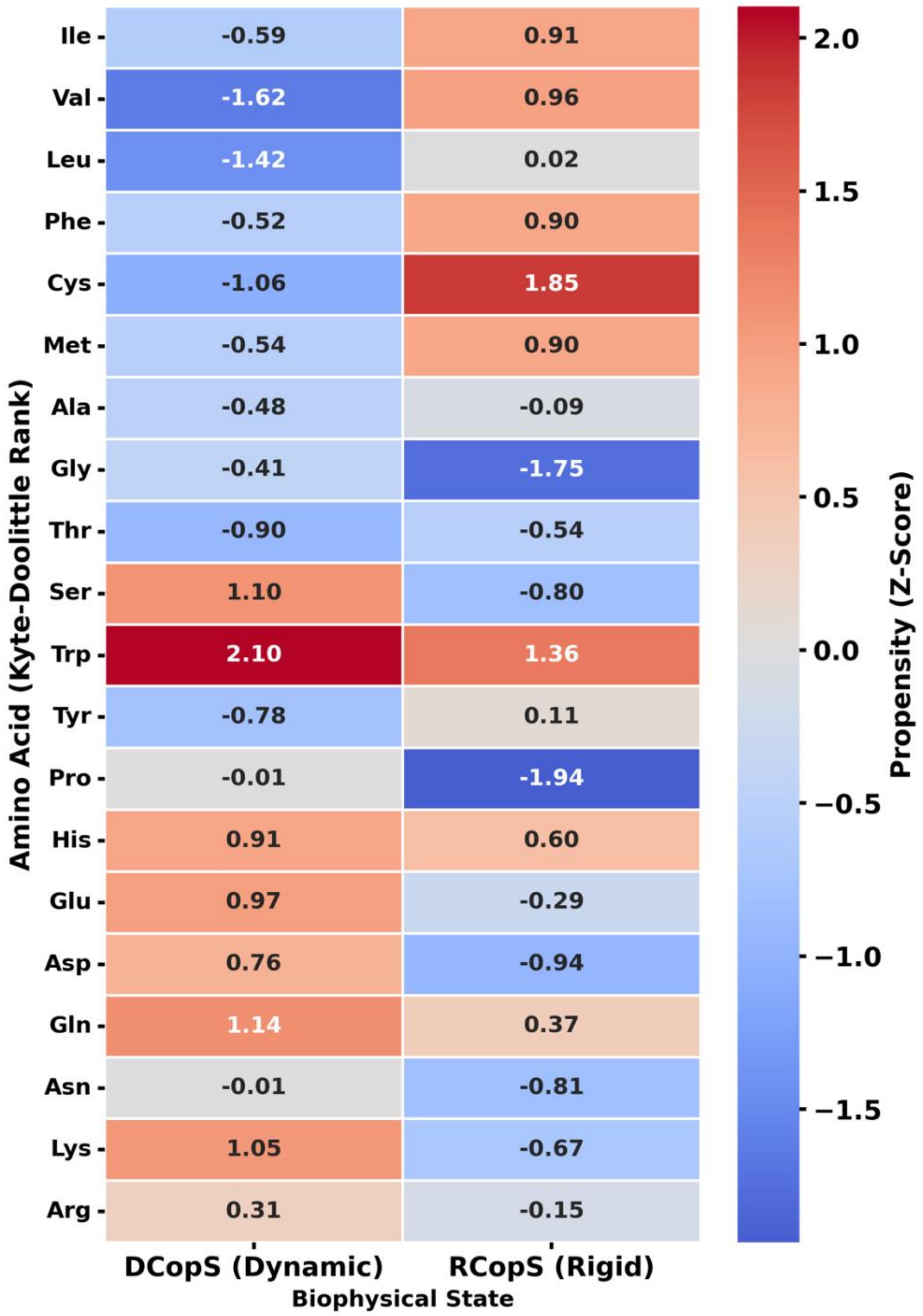


**Figure S11.** Amino Acid Propensity Heatmap of DCopS and RCopS metrics.

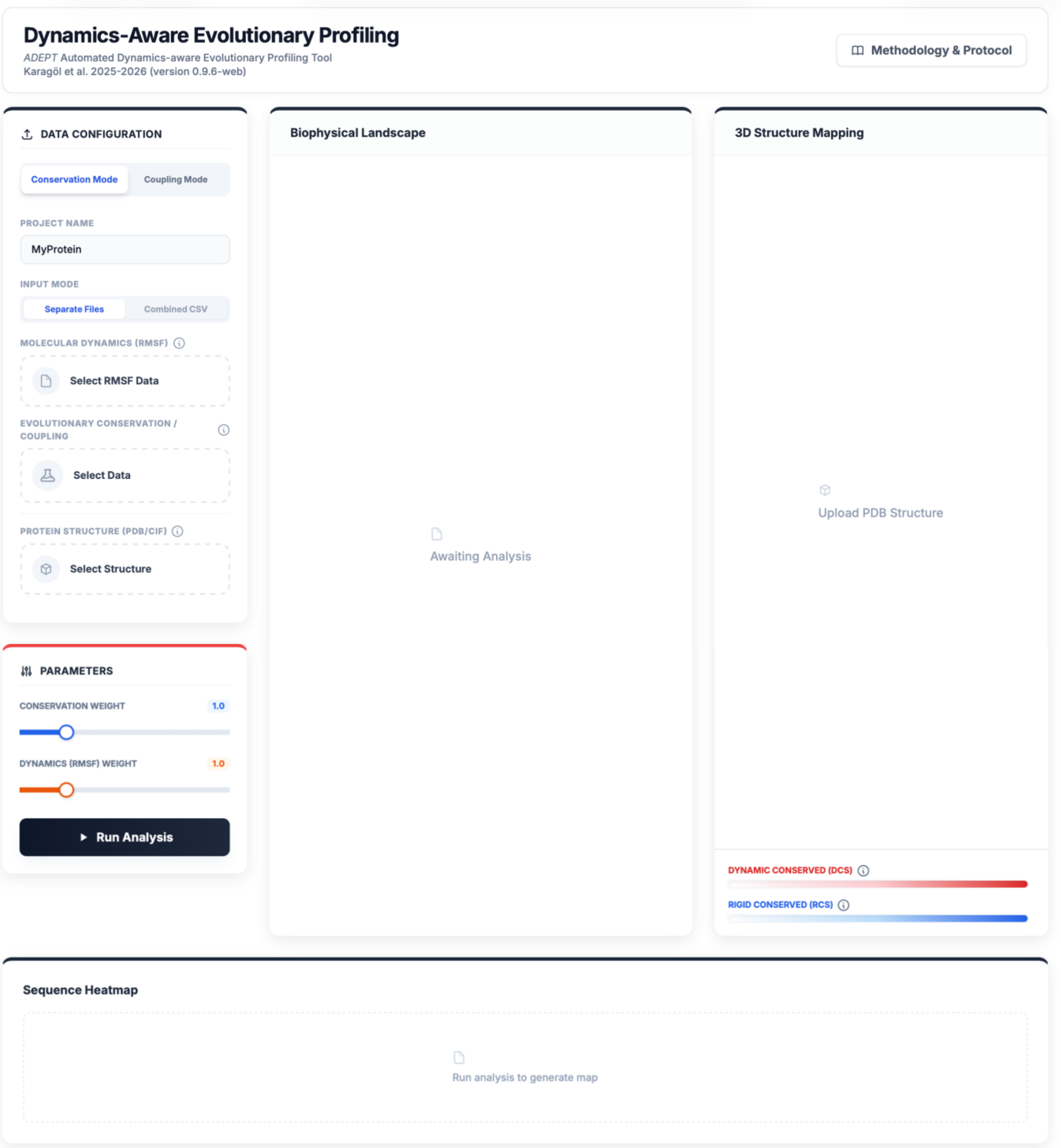


**Figure S12.** Screenshot of *ADEPT* (Automated Dynamics-aware Evolutionary Profiling Tool) web page. (https://www.karagolresearch.com/adept)

**Table S2.** List of available variants from our analysis that are also available in the ClinVar database (likely pathogenic and/or pathogenic variants only).

| Protein_ID | UniProt_ID | Gene_Name | Protein_Change | DCS_Score | RCS_Score | AM_ Score | ClinVar_Significance | ClinVar_Accession |
|---|---|---|---|---|---|---|---|---|
| proteinh46 | O43776 | NARS1 | p.Thr17Met | 5.77521546 | 7.95821059 | 0.0926 | Likely pathogenic | VCV000986308 |
| proteinh46 | O43776 | NARS1 | p.Lys60Thr | 4.00639923 | 7.81799684 | 0.5261 | Likely pathogenic | VCV001700235 |
| proteinh25 | P00558 | PGK1 | p.Lys30Thr | 3.25673338 | 0.62755102 | 0.4744 | Likely pathogenic | VCV000625219 |
| proteinh25 | P00558 | PGK1 | p.Asp315Asn | 2.67962719 | 7.80165816 | 0.9902 | Likely pathogenic | VCV001506855 |
| proteinh3 | P26038 | MSN | p.Asp534Ala | 2.4318675 | 5.62064272 | 0.7441 | Likely pathogenic | VCV000804017 |
| proteinh25 | P00558 | PGK1 | p.Leu88Pro | 2.11855213 | 2.75985128 | 0.9854 | Pathogenic | VCV000009946 |
| proteinh25 | P00558 | PGK1 | p.Thr378Pro | 1.94739253 | 4.21664677 | 0.9588 | Pathogenic | VCV000009956 |
| proteinh25 | P00558 | PGK1 | p.Asp285Val | 1.92554557 | 8.03571429 | 0.988 | Pathogenic | VCV000009951 |
| proteinh25 | P00558 | PGK1 | p.Val266Met | 1.57763655 | 5.38551617 | 0.951 | Pathogenic | VCV000009944 |
| proteinh22 | Q9H7Z6 | KAT8 | p.Lys175Glu | 1.40072916 | 1.26938159 | 0.9993 | Pathogenic | VCV000976462 |
| proteinh22 | Q9H7Z6 | KAT8 | p.Lys175Thr | 1.40072916 | 1.26938159 | 0.9962 | Likely pathogenic | VCV001329928 |
| proteinh25 | P00558 | PGK1 | p.Cys316Arg | 1.3832589 | 5.31089883 | 0.9905 | Pathogenic | VCV000009948 |
| proteinh25 | P00558 | PGK1 | p.Arg206Pro | 1.24916379 | 5.25797872 | 0.8115 | Pathogenic | VCV000009943 |
| proteinh25 | P00558 | PGK1 | p.Ile47Asn | 1.24334798 | 6.28362462 | 0.9973 | Pathogenic | VCV000009952 |
| proteinh25 | P00558 | PGK1 | p.Ser320Asn | 1.08210237 | 6.24430091 | 0.9549 | Pathogenic | VCV000009953 |
| proteinh25 | P00558 | PGK1 | p.Cys50Trp | 0.99629656 | 4.84524262 | 0.9987 | Likely pathogenic | VCV001687119 |
| proteinh25 | P00558 | PGK1 | p.Ser62Asn | 0.99281612 | 6.71550695 | 0.9942 | Likely pathogenic | VCV000391568 |
| proteinh25 | P00558 | PGK1 | p.Asp164Val | 0.98935074 | 9.00911854 | 0.9987 | Pathogenic | VCV000009954 |
| proteinh25 | P00558 | PGK1 | p.Ala354Pro | 0.89768392 | 2.78468302 | 0.9487 | Likely pathogenic | VCV001506865 |
| proteinh25 | P00558 | PGK1 | p.Asp268Asn | 0.87869967 | 1.81393834 | 0.1682 | Pathogenic | VCV000009942 |
| proteinh26 | Q3YBR2 | TBRG1 | p.Ala210Thr | 0.78132944 | 5.20144657 | 0.0969 | Likely pathogenic | VCV000996584 |
| proteinh25 | P00558 | PGK1 | p.Gly158Val | 0.67394095 | 4.78818389 | 0.9952 | Pathogenic | VCV000009947 |
| proteinh52 | Q9H040 | SPRTN | p.Tyr117Cys | 0.53461618 | 6.18179543 | 0.4674 | Pathogenic | VCV000143916 |
| proteinh22 | Q9H7Z6 | KAT8 | p.Met217Leu | 0.50597802 | 4.19456031 | 0.794 | Likely pathogenic | VCV002683826 |
| proteinh24 | P54868 | HMGCS2 | p.Asp136Gly | 0.48261225 | 0.99257963 | 0.9681 | Likely pathogenic | VCV001327468 |

| | | | | | | | | |
|---|---|---|---|---|---|---|---|---|
| proteinh24 | P54868 | HMGCS2 | p.Val141Asp | 0.4373275 | 2.39868044 | 0.9622 | Likely pathogenic | VCV001327463 |
| proteinh22 | Q9H7Z6 | KAT8 | p.Lys181Asn | 0.24001222 | 1.71821534 | 0.9818 | Pathogenic | VCV000976463 |
| proteinh24 | P54868 | HMGCS2 | p.Glu74Lys | 0.22019061 | 2.26038743 | 0.9304 | Likely pathogenic | VCV001327462 |
| proteinh24 | P54868 | HMGCS2 | p.Gly212Arg | 0.2128617 | 8.40290199 | 0.9716 | Pathogenic/Likely pathogenic | VCV000009259 |
| proteinh24 | P54868 | HMGCS2 | p.Gly219Glu | 0.12474414 | 3.39906778 | 0.9653 | Likely pathogenic | VCV003896593 |
| proteinh27 | Q02643 | GHRHR | p.Arg94Trp | 0.08629653 | 2.71380786 | 0.6164 | Likely pathogenic | VCV001449976 |
| proteinh27 | Q02643 | GHRHR | p.Arg94Gln | 0.08629653 | 2.71380786 | 0.2761 | Pathogenic/Likely pathogenic | VCV000161436 |
| proteinh24 | P54868 | HMGCS2 | p.Tyr167Cys | 0.0810042 | 3.31068489 | 0.7319 | Pathogenic | VCV000009262 |
| proteinh24 | P54868 | HMGCS2 | p.Ala171Val | 0.07501354 | 2.45557137 | 0.4387 | Likely pathogenic | VCV000870423 |
| proteinh24 | P54868 | HMGCS2 | p.Gly55Asp | 0.06409203 | 2.36807632 | 0.9764 | Likely pathogenic | VCV002628076 |
| proteinh24 | P54868 | HMGCS2 | p.Phe174Leu | 0.05413154 | 1.28795812 | 0.7558 | Pathogenic/Likely pathogenic | VCV000009257 |
